# Investigating the genomic landscape of mouse models of breast cancer metastasis

**DOI:** 10.1101/2025.01.15.633216

**Authors:** Christina R. Ross, Karol Szczepanek, Jack Sanford, Tinghu Qiu, Larry Pearce, Kent W. Hunter

**Affiliations:** Laboratory of Cancer Biology and Genetics, Center for Cancer Research, National Cancer Institute, NIH; Johnson & Johnson; AstraZeneca; NYU Langone Health

## Abstract

Metastasis remains a major cause of cancer mortality. AbstractThis study, expanding upon previous findings in the MMTV-PyMT model, investigated four independent mouse models, representing luminal (MMTV-PyMT, MMTV-Myc), HER2-amplified (MMTV-Her2) and triple negative (C3(1)TAg) breast cancer subtypes. Consistent with previous results, limited evidence for metastasis-associated somatic point mutations was found for all models. We also found that oncogenic drivers significantly influenced the number and size of metastasis-specific copy number variations (MSCNVs), but common driver-independent MSCNVs were rare. Furthermore, analyzing a cohort with varying genetic backgrounds while maintaining a constant oncogenic driver (PyMT) revealed that genetic background profoundly impacts MSCNVs. Transcriptome analysis demonstrated that oncogenic drivers strongly shaped metastasis-specific gene expression (MSGE), with each driver exhibiting distinct expression profiles. In contrast, MSGE in the PyMT-F1 cohort was more variable across strains. Despite the diversity of MSCNV and MSGE, functional analysis revealed that both mechanisms converge on the modulation of key cellular processes, including immune responses, metabolism, and extracellular matrix interactions. These findings emphasize the complex interplay between oncogenic drivers and genetic background in shaping the genomic and transcriptional landscapes of metastatic lesions.

## Introduction

In the United States, breast cancer is the most frequently diagnosed cancer type and accounts for the third highest number of cancer-associated deaths annually^1^. While patients with localized disease have a 5-year survival rate of 99.6% following tumor resection, the 5-year survival rate drops to 31.9% for patients with distant metastasis^1^. This disparity stems from the inability to effectively treat secondary lesions through either surgical resection or current therapeutic strategies. Metastatic tumors reside in microenvironments and exhibit therapeutic resistances that are distinct from those of the primary tumors, warranting further research and development to create new and more specific therapeutic strategies.

Unfortunately, despite many recent advances, the etiology and mechanisms driving metastasis remain unclear. The conventional model of metastatic progression was first proposed by Peter Nowell in 1976, who hypothesized that tumors arise from a single mutated cell and continue to acquire additional mutations that persist if they enable survival and growth^2^. In this model, clones continue to evolve within the primary tumor (PT), ultimately developing into subclones with metastatic ability. Recent increased access to DNA-sequencing technologies has enabled several clinical studies to characterize the genomic landscape of metastatic tumors^3–5^. However, although much of the data obtained to date is consistent with this hypothesis, no metastasis-driver mutations or acquired genomic aberrations have been identified in any cancer type. Indeed, current data suggest the acquisition of transcriptomic and epigenetic switching mechanisms, rather than mutations, drive dissemination and secondary tumor growth ^6–9^.

To better understand the etiology of breast cancer metastasis, well-credentialed, physiologically relevant, and accessible experimental systems are required. Clinical data, while extraordinarily valuable, is difficult and expensive to obtain in large numbers, experimentally limited, and is usually confounded by treatment, which may significantly influence data interpretation. Mouse models of metastatic cancer therefore provide a complementary experimental vehicle to assess the etiology of metastasis in a treatment-naïve, highly controlled setting. Indeed, several genetically engineered mouse models (GEMMs) have been developed to mimic the molecular subtypes of metastatic breast cancer found in patients. Our laboratory and many others have published extensively using the mammary-specific polyomavirus middle T antigen overexpression mouse model (MMTV-PyMT) to model the luminal B subtype of human breast cancers^6,10–12^. This model was developed in 1992 and faithfully recapitulates human breast cancer progression and metastasis to the lung^13,14^.

Previously, we performed genomic and transcriptomic analysis of the MMTV-PyMT model. By performing whole exome-sequencing (exome-seq) on genomic DNA (gDNA) from matched PT and lung metastatic (lung met) tissues, we showed that very few single-nucleotide variants (SNVs) occur in metastases at a high frequency that are not also found in the primary tumor, and concluded that SNVs are an unlikely driver of metastasis^6^. In addition, we observed no obvious metastasis-associated translocations, but some metastasis-associated copy number variation (CNV). In this study, we expand upon this work to include both the MMTV-Myc and the C3(1)TAg models, as well as provide greater depth to our prior analysis of the MMTV-Her2 and PyMT models^6^. In addition, we have further investigated how an individual’s personal ancestry might influence metastasis-associated genomic evolution in the MMTV-PyMT model by analyzing F1 animals generated from crosses between phylogenetically related mouse strains that exhibit significant differences in metastatic efficiency. Consistent with previous analyses, we confirm that SNVs with a metastasis driver function are not present in most of the animals analyzed, regardless of oncogenic driver or genetic background. However, we have identified genetic background as a strong determinant of PT stability, copy number region size, and gene-set enrichment within regions of gDNA gain or loss. Further, we show that gene expression at end-stage is not strongly correlated with CNV in our models. Finally, our work reveals no common gene or gene-set found within CNV, but rather cellular processes enriched in metastasis-specific gene expression and alternative splicing.

## Results

### Single nucleotide variants are unlikely drivers of metastasis

To evaluate the relative contributions of oncogenic driver and genetic background to the genomic landscape of primary tumors and lung metastases, we developed two mouse model cohorts (Fig 1a). The first cohort, the “oncogenic driver” cohort, utilized four distinct mouse models of metastatic breast cancer, each with a unique oncogenic driver but identical FVB genetic background. The mouse mammary tumor virus (MMTV)-neu (Her2) line was used to model Her2-amplified (Her2+) breast cancer, a subtype representing approximately (17%) of breast cancer patients^15–17^. The MMTV-Myc (Myc) line was used to model metastatic Myc-driven breast cancer, which is observed in approximately 30-50% of high-grade tumors^18^. The C3(1)/TAg-REAR (C3Tag) line was used to model a metastatic basal-like subtype, which is seen in 8% of breast cancer patients^19,20^. Finally, the N-Tg(MMTV-PyVT)634Mul/J (PyMT) line, which forms tumors that are highly metastatic, was used to model activated tyrosine kinase-driven cancers, which represent 49% of Luminal B, 32% of Luminal A, 42% of Her+, and 7% of Basal-like tumors, or 40% of all breast cancers^21^. Thus, the “oncogenic driver” cohort was comprised of four groups, PyMT, Her2, C3Tag, and Myc, with the FVB genetic background held constant. For the second cohort, or “PyMT-F1” cohort, FVB-PyMT males were bred to females from seven different genetic backgrounds and tumor bearing PyMT-F1 offspring were evaluated. The genetic backgrounds chosen constitute two pairs (MOLF and CAST, BL6 and BL10) and one group of three (AKR, BALB, SEAGN) closely related strains with inverse metastatic efficiency (high(H) or low (L)) (Fig 1a and Table S1)^12^, permitting us to evaluate the contribution of genetic background on the genomic landscape with oncogenic driver held constant. We isolated genomic DNA (gDNA) from matched pairs of primary tumors (PT) and lung metastases (Lung mets) from both cohorts (Table S1) and performed exome-sequencing (exome-seq) (Fig 1a).

**Figure 1.**
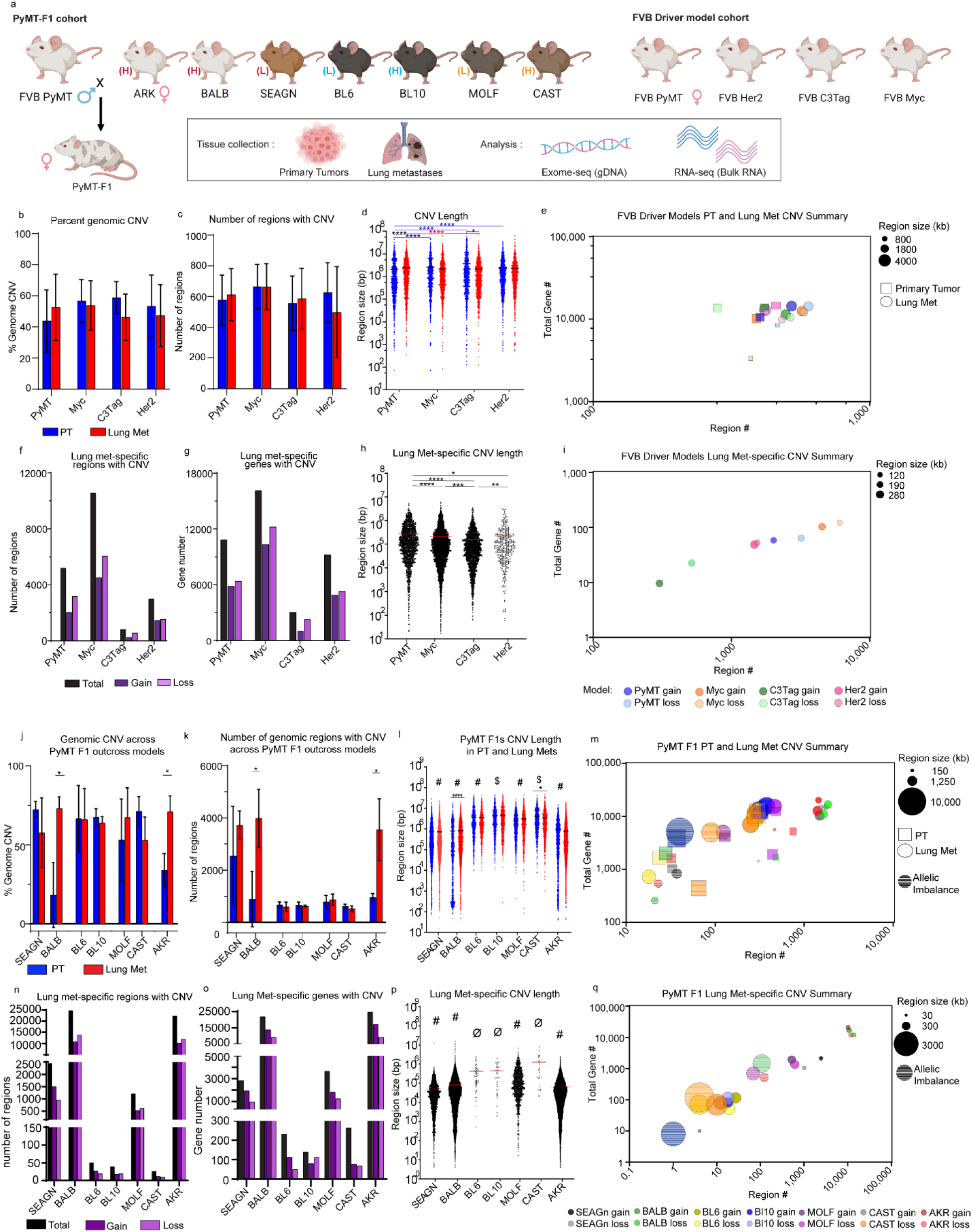
Model and genetic background shape MSCNV (a) Schematic overview of mouse models, tissues collected, and analyses performed in this study. (b-c) Histograms showing (b) the average percentage of the genome and (c) the average number of regions with CNV with standard deviation (SD), PT (blue) and lung mets (red). (d) Dot plot of individual CNV region lengths from aggregate data (the average indicated by black line). (e) Multivariate bubble plot showing average CNV number and aggregate total number of genes with CNV for PT (squares) and lung met (circles) from each model, with bubble size corresponding to CNV region size. (f) Total number (black) of regions (f) and genes (g) with significant CNV gain (dark purple) and loss (light purple) in MSCNV. (h) Dot plot showing individual CNV region lengths from MSCNV (the average indicated by red line). (i) Multivariate bubble plot showing average CNV number and aggregate total number of genes with CNV for MSCNV (circles) from each model, with bubble size corresponding to CNV region size. (j) Histogram showing the average % of the genome and (k) and average number or regions with CNV with SD, PT (blue) and lung mets (red). (l) Dot plot showing individual CNV region lengths from aggregate data with all animals per model combined (the average indicated by a line). (m) Multivariate bubble plot showing average CNV number and aggregate total number of genes with CNV for PT (squares) and lung met (circles) from each model, with bubble size corresponding to CNV region size. (n) Total number (black) of regions and (o) genes with significant CNV gain (dark purple) and loss (light purple) in Lung mets compared to PT. (p) Dot plot showing individual CNV region lengths from MSCNV (the average indicated by red line). (q) Multivariate bubble plot showing average CNV number and aggregate total number of genes with CNV for MSCNV (circles) from each model, with bubble size corresponding to CNV region size. Significance determined by Kruskal-Wallis test (d, h, l, p) or unpaired t-test (b, c, j, k). *(p<0.05), **(p<0.01), ***(p<0.001) ****(p<0.0001), $ = significantly different from 6 groups, # = significantly different from 5 groups, ø = significantly different from 4 groups.

We have previously shown that high allele frequency, metastasis-specific SNVs were not found at consequential rates in a smaller mouse cohort^6^. Using the exome-seq data from this larger cohort described above, we expanded the number of individuals used for SNV analysis. Consistent with our previous data, high frequency, metastasis-specific SNVs were not observed, suggesting that common SNVs do not function as major drivers of metastasis (Fig S1). Therefore, subsequent analysis focused on copy number variation (CNV) induced by regions of genomic gain or loss. Herein, through use of Nexus Copy Number Software 10.0, we have evaluated CNV differences across sites (PT or Lung mets), both within and between cohorts.

### Oncogenic driver has little influence over CNV event number

We first evaluated the oncogenic driver cohort, visualizing PT and LM data separately for each FVB model, and calculated the average percent of the genome with CNV (Table S2). We observed no significant difference in the percentage of the genome with DNA gain or loss between PT and Lung mets, and no difference between oncogenic driver models (Fig 1b). The average percentage of the genome with CNV ranged between 43-58% in PTs and 46-53% in Lung mets (Fig 1b and Fig S2). Similarly, the average number of regions with CNV did not differ significantly between tissue type or model (Fig 1c and Table S2).

Despite having a similar degree of CNV overall, we observed key differences in the size of DNA regions with amplification or deletion. While no significant differences were noted in the CNV length for the Myc and Her2 models, CNV regions in the primary tumors were shorter in PyMT and longer in C3Tag PTs compared to their matched Lung mets (Fig 1d and Table S3). When analyzing regions of gain or loss separately, we found that regions of DNA gain were significantly larger (3572 – 4057 kb) than those of loss (806 – 1340 kb) in PTs from three models (Her2, Myc, and PyMT), and in Lung mets from all models (2497 – 3483 kb gains, 1370 – 2330 kb losses) (Fig S3a-b). However, despite these slight but consistent differences in CNV gain and loss length, the four FVB oncogenic driver models maintain a similar number of regions and genes impacted by CNV (Fig 1e).

### Oncogenic driver determines metastasis-specific CNV event number and size

We next focused on metastasis-specific CNV (MSCNV) for each model, or regions altered in Lung mets but not PTs. Interestingly, oncogenic driver played a significant role in the number of MSCNV regions and the number of genes within them (Table S4). The Myc driven model had the highest number of MSCNV regions (10,576) impacting the highest number of genes (16,134), followed by PyMT (5,198 regions, 10,823 genes), Her2 (3,003 regions, 9,213 genes), and C3Tag (818 regions, 3007 genes) (Fig 1f-g and Table S4 and Table S5). Regions of MSCNV were even shorter in length than the average Lung met CNV by around 10-fold, 200 kb compared to 2000 kb, respectively (Fig 1h and Fig Sc). Overall, there were small but significant differences in the average length of MSCNV between models (Fig 1h). When evaluating CNV gain and loss separately, we again observed significant differences in the length of MSCNV regions in a model-dependent manner (Fig S3c and Table S4). Myc and C3TAg MSCNV regions of gain were significantly longer than regions of loss, while the opposite was observed for PyMT, and no difference was observed for Her2. This data reveals an oncogenic driver-dependent influence on MSCNV number and size, thus impacting the number of genes with Lung met-specific CNV (Fig 1i).

### Genetic background alters CNV event number and size

We next performed similar analyses on samples from our PyMT-F1 cohort to evaluate the contribution of genetic background to CNV when the oncogenic driver (PyMT) was held constant (Fig 1a and Table S1). While the percentage of the genome impacted by CNV was similar in Lung met tissue across strains (53-73%), significantly less was observed in PTs from BALB and AKR F1 mice, at only 18.24% and 33.89%, respectively (Fig 1j and Fig S4a and Table S2). More specifically, BALB and AKR had fewer regions of CNV within PT gDNA compared to matched Lung met tissue (Fig 1k). Interestingly, when comparing the number of regions of Lung met CNV between all F1s we noted a divergence where gDNA from BL6, BL10, MOLF, and CAST tissues had between 600-900 regions, but gDNA from BALB, SEAGN and AKR had 3000 to 4000 regions (Fig 1k). Due to the heterozygous genetic background of the F1 PyMT mice, we were also able to assess regions of allelic imbalance. While gDNA from most strains had a similar number of regions with allelic imbalance in PT and Lung mets (18-40), CAST and MOLF, closely related wild-derived strains, had a much higher number (65-128) (Fig S3d and Fig S5 and Table S2).

When measuring CNV region size in tissue from PyMT F1s we observed an inverse relationship with region number. BL6, BL10, MOLF, and CAST F1 mice had larger regions of CNV than SEAGN, BALB, and AKR (Fig 1l and Table S3). We theorized that this inverse relationship was likely responsible for the relatively constant percentage of the genome impacted by CNV, excluding BALB and AKR PTs. In addition to differences between strains, we also observed small but significant differences in the length of CNVs in lung mets compared to PTs in CAST and BALB F1s (Fig 1l). Analysis of region size was further broken down to genomic gains, losses, and allelic imbalance (Fig S3e-f). Regions of genomic loss trended larger than those of gain in both PT and Lung mets, regardless of background (Fig S32-f). Further, in some F1 strains the regions of allelic imbalance in PT and Lung met gDNA were significantly larger than the average CNV size. In strains with larger allelic imbalance, the size was roughly double the length of average CNV (Fig S3e-f and Table S3). Taken together, PyMT F1 models have unique CNV fingerprints shaped by CNV region number and region size, therefore altering the number of genes impacted by CNV (Fig 1m).

### Genetic background alters metastasis-specific CNV event number and size

We next focused on MSCNV among the PyMT F1 cohort. First, the closely related BL6 and BL10 strain pair had the fewest number of MSCNV, with only 50 and 39, respectively (Fig 1n and Table S4). Closely related BALB, SEAGN, and AKR had the highest number of MSCNV, ranging from 2,448-24,650. Though both MOLF and CAST are wild-derived strains, they exhibited vastly different numbers of MSCNV regions, at 1,201 and 26, respectively. Interestingly, the number of regions of allelic imbalance within MSCNV did not appear to correlate with strain relatedness and ranged from a single region observed in BL10 F1s to 121 regions in AKR F1s (Fig S3g).

As we observed with the oncogenic driver models (Fig 1g), the number of genes impacted by MSCNV correlated closely with the number of regions, rather than the size of CNV region. BL6, BL10, and CAST had the smallest number of genes impacted by CNV, ranging from 139-264 (Fig 1o and Table S5). BALB, SEAGN, and AKR had the highest number of genes contained within MSCNV, ranging from 2820-24,615 (Fig 1n). Further, the number of genes altered by CNV appeared to be evenly distributed between regions of gain and loss. Finally, the number of genes found within regions of metastasis-specific allelic imbalance also correlated with the number of regions rather than region size (Fig S3g-i).

MSCNV region size again followed an inverse relationship with region number. SEAGN, BALB, and AKR had the smallest regions of MSCNV, ranging from 49-83 kb, whereas CAST had the largest average size of 1,250 kb (Fig 1p and Table S4). Comparison of MSCNV length between F1 strains revealed that each strain had a statistically different MSCNV length from all other F1s, and only BL6 and BL10 were not statistically different from one another. When region size for MSCNV was further broken down to gDNA gain, loss, and allelic imbalance, we noted once again that regions of allelic imbalance were larger than gain or loss for six of the seven F1 strains, a difference that was statistically significant in BALB and AKR (Fig S3i). Metastasis-specific regions of gain and loss varied greatly in size across F1 models, but there was little difference in size when comparing regions of loss and gain within the same F1 strain (Fig S3i). Together, this data reveals that metastasis-specific CNV follows a linear trend between CNV region number and region length, as well as between region number and the number of genes with CNV, which is modulated in a strain-dependent manner (Fig 1q).

### Genetic background shapes gene set enrichment in CNV

We next determined if specific genes or pathways were commonly enriched in MSCNV. For the FVB oncogenic driver models, 1676 genes were commonly found within regions of MSCNV across all four models, which accounted for 10-56% of the overall number of genes within MSCNV in each model (Fig 2a and Fig S6a). Gene ontology (GO) analysis of this common gene set revealed enrichment of cell adhesion, granzyme mediated cytolysis, and T-cell viral response pathways (Fig S6b and Table S6). Comparison of GO analysis performed with the MSCNV gene sets from each FVB model individually revealed 25 pathways commonly impacted by MSCNV: 11 immune-related, four metabolic, two developmental, and eight uncategorized (Fig 2b, Fig S6c, Table S7). Notably, one of the uncategorized pathways was *Cellular response to nicotine*, which we previously identified in a prior large transcriptomic study and showed that exposure to nicotine significantly increased mammary cancer metastasis in a mouse allograft model^11^.

**Figure 2.**
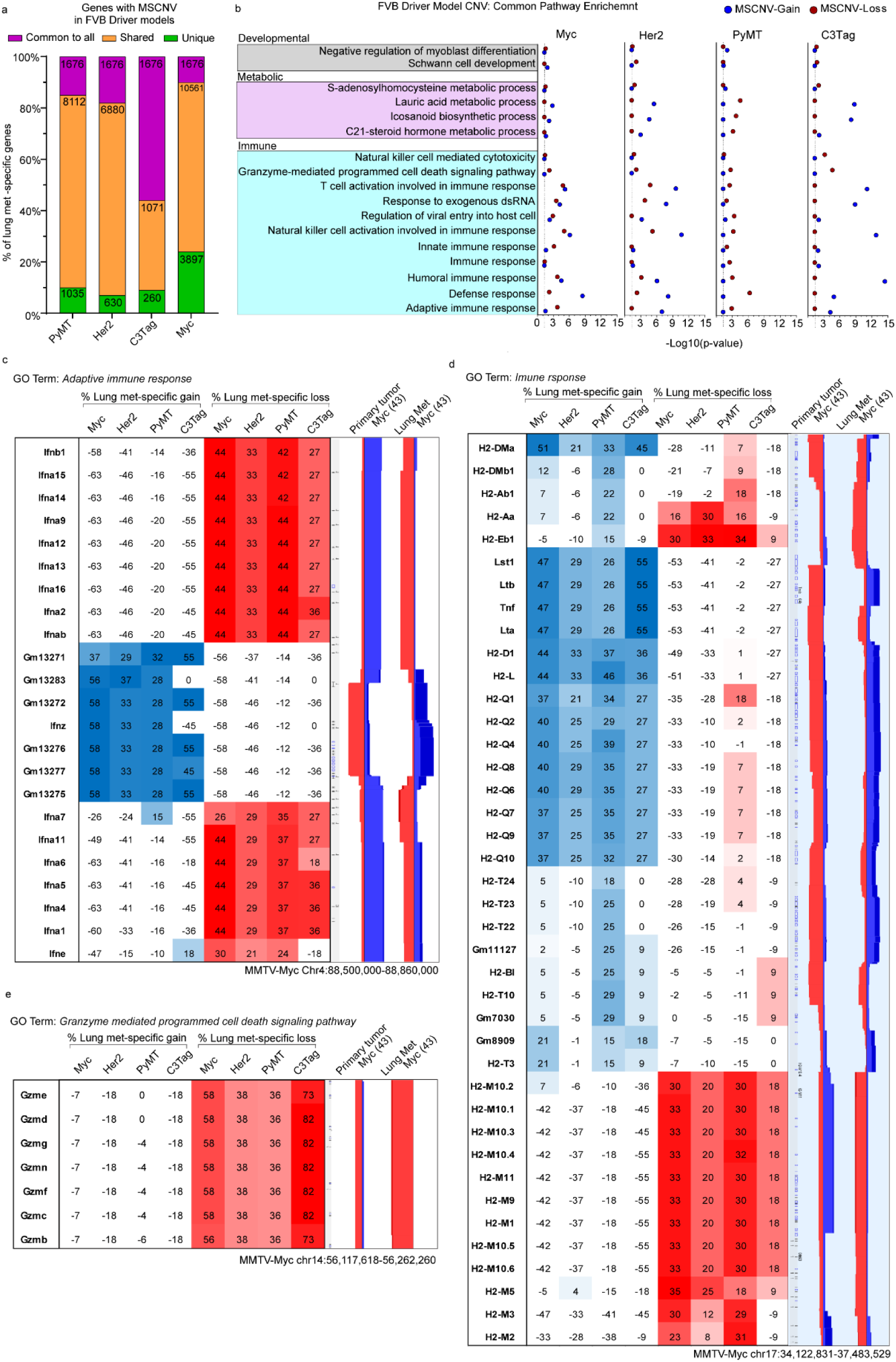
FVB driver models have metastasis-specific regions and gene families with CNV (a) Histogram showing the percent overlap in MSCNV gene lists from each model. Numbers within bars indicate gene number common to all (purple), shared among some (yellow), or unique to each model (green). (b) DAVID analysis clustering of 1676 MSCNV genes common to all models with enrichment in CNV gain (blue) or loss (red) indicated by -Log10(p-value) (dotted line indicates cutoff for significance (≥ 1.3). (c-e) Heatmap indicating % lung met-specific gain in blue (% PT gain - % LM gain) and lung met-specific loss in red (% PT loss - % LM loss) for genes within a single genomic region from the (c) Adaptive immune response, (d) immune response, and (e) Granzyme mediated programmed cell death signaling pathway. CNV signal traces from PT and lung mets aligned using Nexus 10 genomic view and the Myc model data set.

Interestingly, several of the immune-related pathways contained CNV-altered gene families clustered within a single locus. Furthermore, pathways enriched by CNV had both significant metastasis-specific gain and loss (Fig 2b). A 400 kb region of chromosome 4 was altered in six of the 11 immune-related pathways common to the oncogenic driver cohort MSCNV. Within this region, metastatic samples had genomic gain flanked on either side by genomic loss, but the inverse was observed in PTs (Fig 2c). This locus contains a cluster of 23 interferon genes similar to a region housing several ortholog interferon genes on human chromosome 9 p21.3. Additionally, we identified a 3000 kb region on chromosome 17 with both metastasis-specific gain and loss. Within this locus are the 32 H2 genes that make up the major histocompatibility complex (Mhc). Specifically, we observed a common amplification of H2-Q genes and deletion of H2-M genes in metastases and the inverse in PTs (Fig 2d). A third region of interest was observed on chromosome 14 that stretched approximately 150 kb and included seven granzyme (Gzm) genes that can be found within two of the common immune pathways. We observed significant Lung met-specific deletion of this locus across all four FVB driver models (Fig 2e). While several more loci were identified as major contributors to each enriched gene ontology, we noted that PyMT and Myc often aligned but C3Tag and Her2 samples were more variable (Table S8).

We performed a similar analysis for the PyMT-F1 cohort. The 25 common pathways identified from the FVB oncogenic driver model analysis were compared to GO terms enriched in MSCNV from each F1 model individually using several two-group Venn diagram analyses. AKR had the highest overlap of 17 pathways, followed by BALB (6), SEAGN (2) and MOLF (1), while BL6, BL10, and CAST had no overlap (data not shown), suggesting that genes within regions of MSCNV are enriched in a genetic background-dependent manner, as F1s with the highest GO overlap were also the strains most closely related to FVB. Additionally, comparative analysis of gene lists from regions of MSCNV revealed that, except for in BALB F1s, most genes were subject to CNV in more than one strain but no genes were commonly altered in all strains (Fig 3a and Fig S6d and Table S5). Similarly, comparison of gene ontologies enriched by CNV were largely unique to each PyMT-F1 strain and there were no GO terms enriched by MSCNV across all strains (Fig 3b and Fig S6e and Table S9).

**Figure 3.**
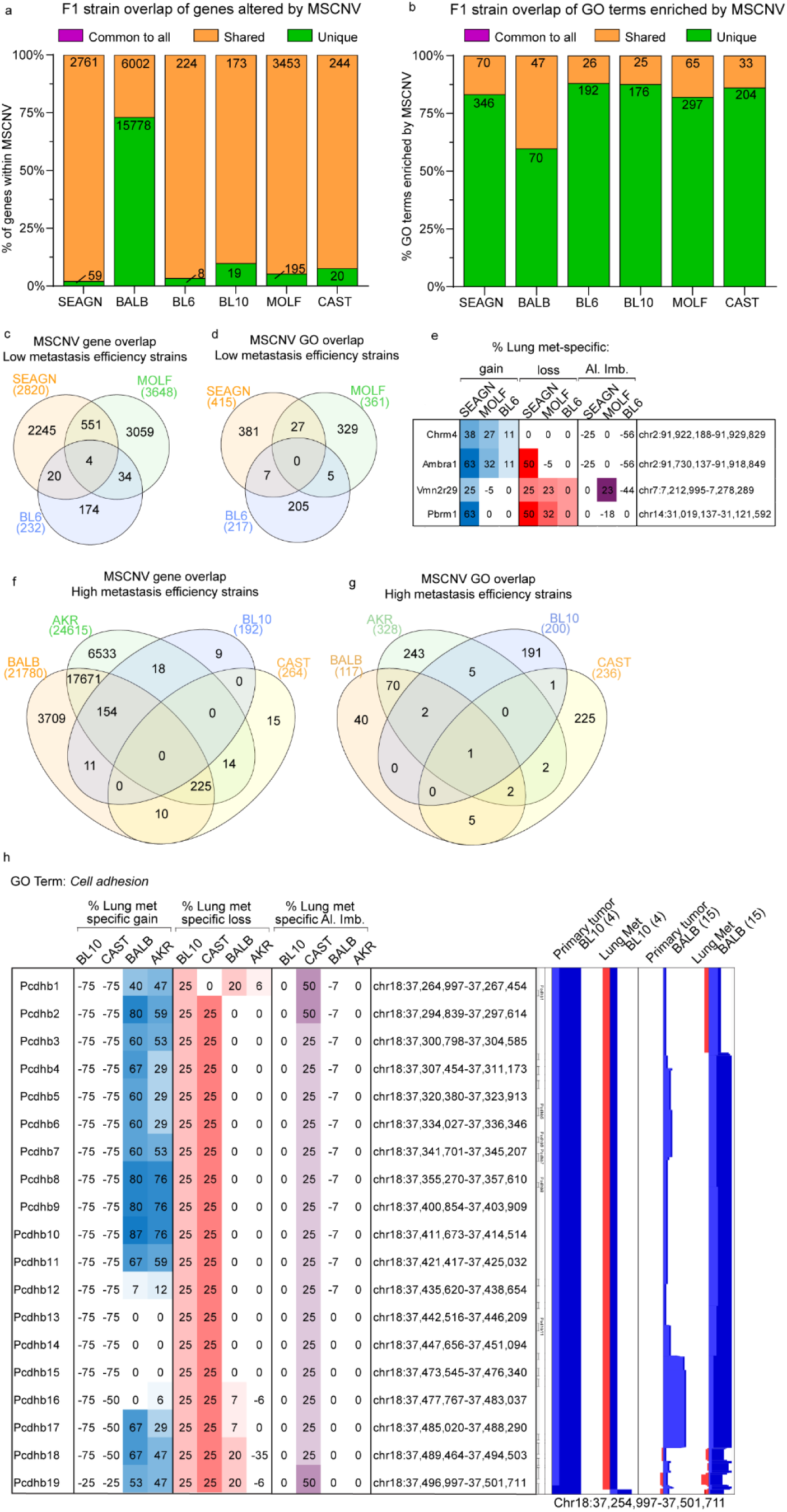
PyMT F1 models have no common gene sets or ontologies enriched by MSCNV (a-b) Histograms of the percent overlap in gene lists (a) and GO terms (b) enriched by MSCNV in each model. Numbers within bars indicate gene number common to all (purple), shared among some (yellow), or unique to each model (green). (c-d) Venn diagram analysis of gene sets (c) and GO terms (d) enriched by MSCNV in low metastatic efficiency strains. (e) Heatmap indicating % lung met-specific gain in blue (% PT gain - % LM gain) and lung met-specific loss in red (% PT loss - % LM loss) for genes with MSCNV in low metastatic efficiency strains. (f-g) Venn diagram analysis of gene sets (f) and GO terms (g) enriched in MSCNV from high metastatic efficiency strains. (h) Heatmap indicating % lung met-specific gain in blue (% PT gain - % LM gain), lung met-specific loss in red (% PT loss - % LM loss), and lung met-specific allelic imbalance (Al. Imb.) in purple (% PT Al. Imb. - % LM Al. Imb.) for genes with MSCNV in high metastatic efficiency strains from the Cell adhesion pathway within a single locus. CNV signal traces from PT and Lung mets aligned using Nexus 10 genomic view of the BL10 and BALB model data sets.

We previously reported differing metastatic efficiency of the mouse strains included in this study (Fig 1a and Table S1)^12^. Therefore, we next asked if F1-PyMT metastases from strains with either high or low metastatic efficiency might have similar genomic evolution and common MSCNV. To test this, genes within CNV gain, loss, and allelic imbalance, as well as enriched GO terms were compared among strains with low (SEAGN, BL6, and MOLF) and high (BALB, AKR, BL10, and CAST) metastatic efficiency. For low metastatic efficiency strains, we identified four genes commonly altered by MSCNV, but no commonly enriched GO terms (Fig 3c-d). The *Chrm4* and *Amrba1* genes are found within a region of metastasis-specific genomic gain on mouse chromosome 2 (human chr11 p11.2), whereas *Vmn2r29* and *Pbrm1* are found on mouse chromosome 7 and 14, respectively, and have MSCNV loss (Fig 3e). In contrast, high metastatic strains revealed no common genes within regions of MSCNV, but all strains had enrichment of the *Cell adhesion* GO term (Fig 3f-g). Heatmap analysis showed high variability between strains for genomic gain or loss of factors within the *Cell adhesion* ontology gene list, except for a locus on mouse chr18 encompassing a stretch of 19 protocadherin genes (Fig 3h and Table S4). BALB and AKR showed amplification of this region in metastatic tissue compared to PTs, whereas BL10 and CAST showed amplification within the PT DNA not seen in the metastases, therefore resulting in a comparative metastasis-specific loss in this region (Fig 3h).

As well as genes within regions of loss and gain, we assessed imbalance between the FVB and non-FVB allele induced by CNV and enriched in Lung mets. Allelic imbalance of a common gene or gene set was not observed across this cohort (Fig S7a). Additionally, we did not observe allelic imbalance common to strains with either high or low metastatic efficiency (Fig S7b-c). However, when we observed allelic imbalance within each closely related strain pair, common regions of imbalance were identified. In BALB and SEAGN PyMT F1 mice we observed common allelic imbalance in two regions, Chr1qH3 and Chr11qB4 (Fig S7d). Interestingly, four genes found within a region of amplification on Chr1qH3 had an allelic imbalance in BALB Lung mets that was not observed in the PTs, but SEAGN showed the inverse with allelic imbalance in the PT not observed in Lung mets. The second region encompassed three genes with allelic imbalance in both BALB and SEAGN PTs that was not enriched in Lung mets (Fig S7d). These seven genes are similarly arranged within the human genome at loci Chr1 q23.3 and Chr17 p13.2, respectively. MOLF and CAST PyMT F1s also had two common regions of metastasis-specific allelic imbalance. A region of PT and Lung met genomic gain on Chr7q resulted in allelic imbalance in Lung mets from both strains (Fig S7e). This locus houses 11 genes and corresponds to the human Chr10q26 which contains the same set of ortholog genes. A second region within mouse Chr18q was also a site of genomic gain and common Lung met-specific allelic imbalance, encompassing 10 genes that are also found together within human Chr5q31.3 (Fig S7e). The third strain pair, BL6 and BL10, both had metastasis-specific allelic imbalance, but no common regions could be identified (Fig S7f). Taken together, this data suggests that genetic background is a stronger determinant than oncogenic driver in shaping the enrichment of CNV-altered genes in the metastatic genome.

### Metastatic gene expression Is not shaped by CNV

We next asked if genes in MSCNV regions were differentially expressed in the metastatic lesions by performing bulk RNA-sequencing (RNA-seq) of RNA isolated from matched PT and Lung met tissue collected from the oncogenic driver and PyMT-F1 cohorts. We found significant variation in the number of genes within regions of CNV that were expressed in metastatic tumors (Fig 4a and Table S10). CAST had the lowest expression with only 8% of MSCNV genes found in the RNA-seq signal, but BALB had the highest with 58%. We next performed differential gene expression analysis between Lung mets and PTs for each model and asked if genes with MSCNV had altered expression compared to PTs (Table S11). Again, we observed high variability within the data, with the FVB-MYC model showing the lowest number of MSCNV genes with altered expression (0.01%), and PyMT F1-AKR showing the highest (33%). When we examined the genes within MSCNV that also had significant differential gene expression compared to the PT, we noted that the expression change did not correspond to the CNV type (gain or loss) (Fig 4b-c). Taken together this shows a poor association between CNV type and direction of any changes in gene expression at the macro metastasis stage (Fig 4d-n).

**Figure 4.**
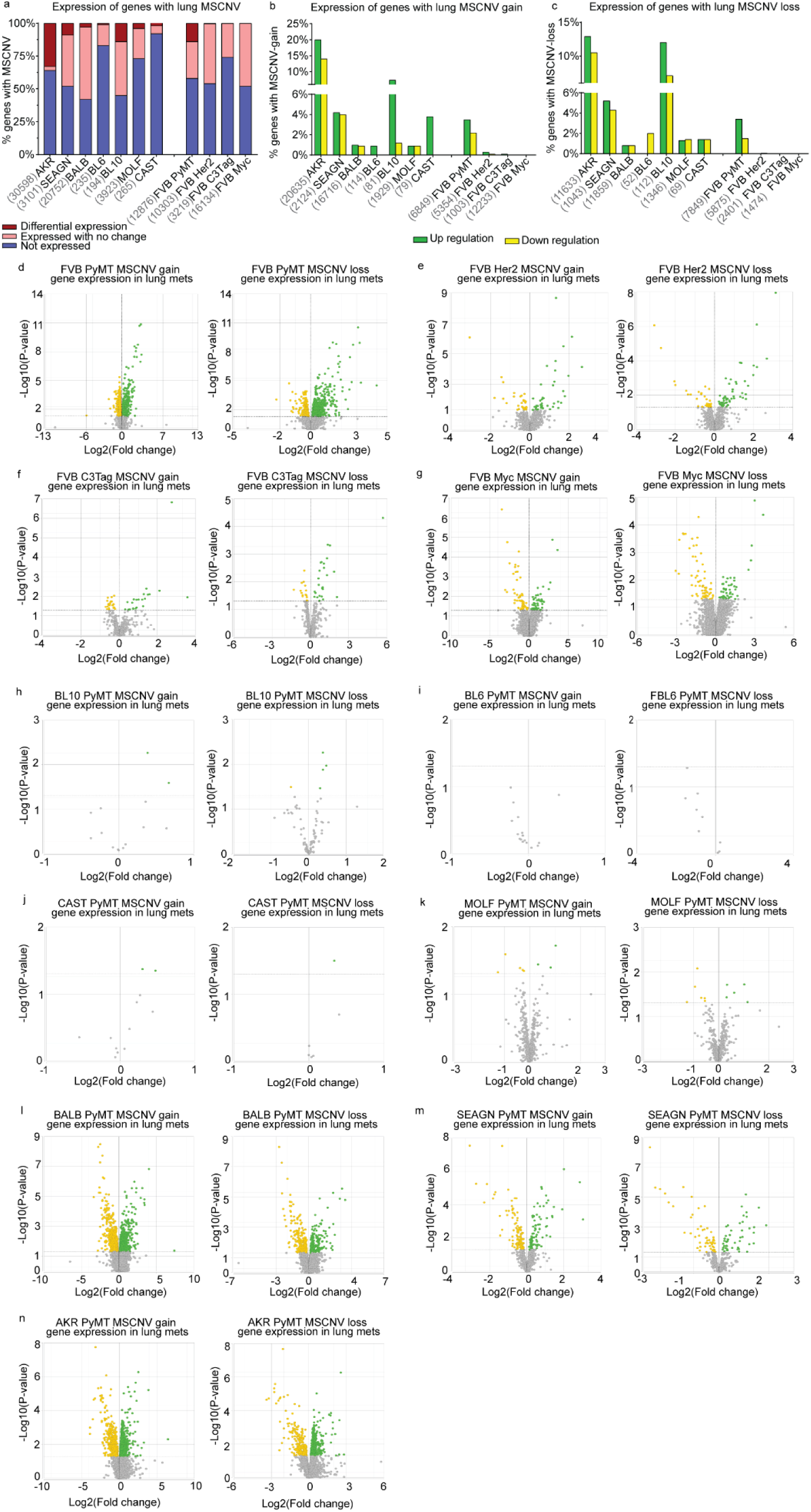
MSCNV does not significantly impact metastasis-specific gene expression (a) Histogram of the percentage of genes with MSCNV that are not expressed (blue), expressed in lung mets with no change from PT (pink), and differentially expressed in lung mets compared to PTs (red). Numbers listed before each model name represent the MSCNV gene set size. (b-c) Histograms showing the percentage of genes with MSCNV-gain (b) or MSCNV-loss (c) that had significant up regulation (green) or down regulation (yellow) in lung mets compared to PTs. (d-n) Paired volcano plots for the MSGE of genes within regions of MSCNV gain or loss for each FVB driver model (d-g) and PyMT-F1 model (h-n).

### Oncogenic driver and genetic background shape metastatic gene expression

As MSCNV was not a major contributor to metastasis-specific gene expression (MSGE) within metastatic lesions, we analyzed our RNA-seq data to determine how oncogenic driver or genetic background impacted MSGE. To begin, all samples were visualized together by unsupervised principal component analysis (PCA), which resulted in four distinct clusters differentiated by oncogenic driver (Fig 5a). When the analysis was limited to individual models within the oncogenic driver cohort there was no significant separation of PT and Lung met samples (Fig 5b-e). Together, these results indicate that oncogenic driver is a stronger determinant of sample similarity than tissue type or genetic background.

**Figure 5.**
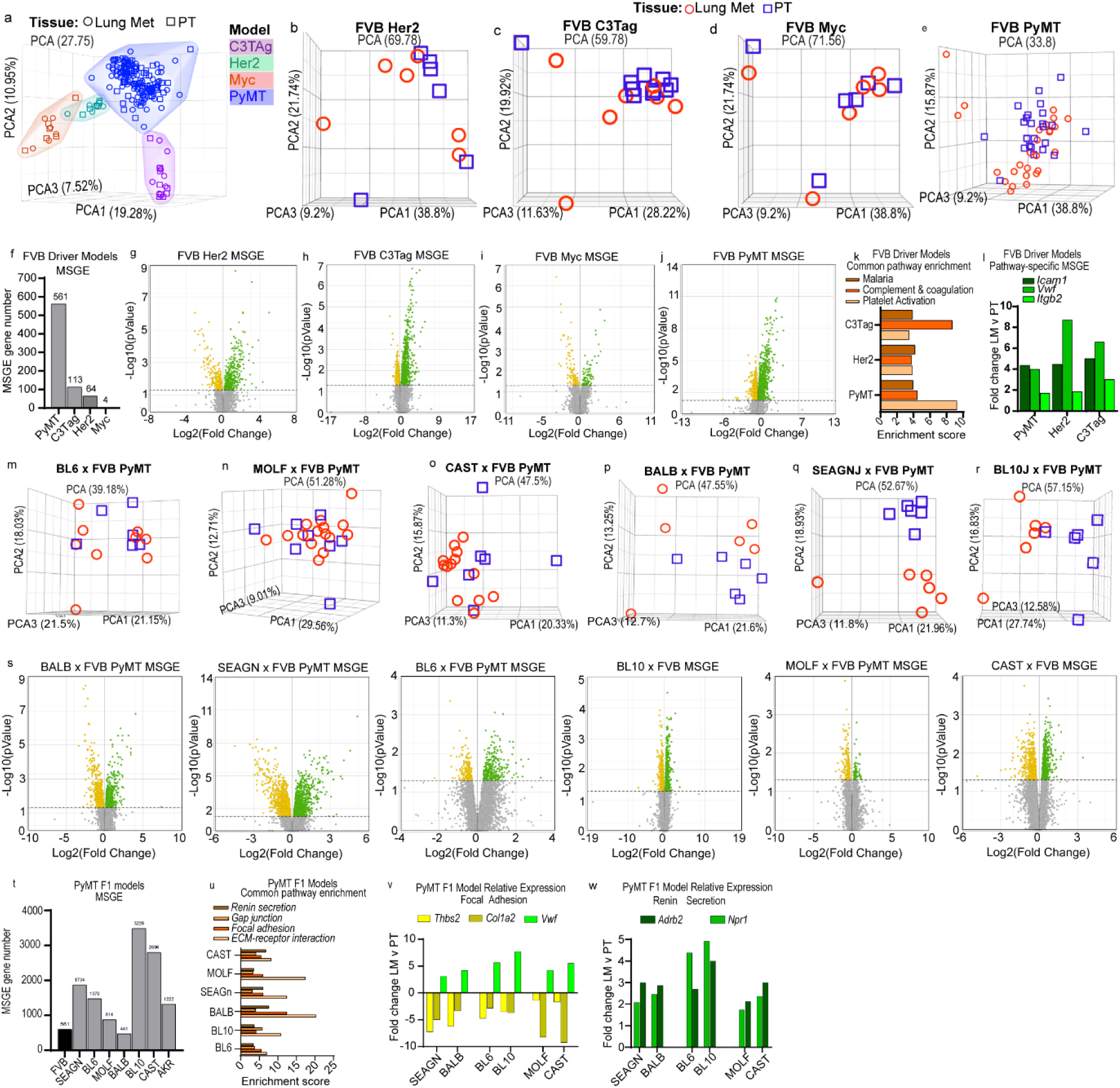
Metastatic gene expression is shaped by oncogenic driver and genetic background. (a) Unsupervised PCA of RNA-seq data from PT and Lung met tissues across all FVB driver and PyMT-F1 mouse models. (b-e) Unsupervised PCA of individual PT and lung met samples from FVB Her2 (b), FVB C3Tag (c), FVB Myc (d), and FVB PyMT (e). (f) Histogram of the number of alternatively expressed genes in lung mets vs. PT for each FVB driver model (FDR<0.05). (g-j) Volcano plots with significantly up regulated genes in green and down regulated genes in yellow for MSGE in FVB Her2 (g), FVB C3Tag (h), FVB Myc (i), and FVB PyMT (j). (k) Histogram showing the enrichment score for commonly enriched pathways from MSGE of the C3Tag, Her2, and PyMT models. (l) Histogram showing the fold change of genes common to pathways enriched by MSGE. (m-r) Unsupervised PCA of individual PT and lung met samples from BL6 (m), MOLF (n), CAST (o), BALB (p), SEAGN (q), and BL10 (r). (s) Volcano plots with significantly up regulated genes in green and down regulated genes in yellow for MSGE in the PyMT-F1 models. (t) Histogram of the number of alternatively expressed genes in lung mets vs. PT for each PyMT-F1 model (FDR<0.05). (u) Histogram showing the enrichment score for commonly enriched pathways from MSGE of the F1-PyMT models. (v-w) Histograms showing the fold change of common MSGE enriched in the Focal Adhesion (v) and Renin Secretion pathways (w).

Gene expression analysis comparing Lung met and PTs revealed that the extent to which MSGE differs from the primary tumor is also oncogenic driver-dependent. Using a cut-off of p<0.05 and FDR<0.1, the PyMT model had significant changes in the expression of 561 genes, C3Tag 113, Her2 64, and Myc only 4 genes (Fig 5f and Table S11). While the Her2 and Myc models showed an even distribution of up- and down-regulated genes, the PyMT and C3Tag models favored up-regulation of metastasis-specific genes (Fig 5g-j). Alternatively regulated genes in the C3Tag and Her2 models overlapped with those from PyMT, but only a single gene, *Pcdh17*, was commonly altered in metastatic tissue from all four models (Fig S8a-b). According to The Genotype-Tissue Expression (GTEx) project, *PCDH17* is highly and specifically expressed in Lung tissue and therefore may simply represent the stroma in which the tumor cells reside (Fig S8c). However, this is challenged by the observation that *Pcdh17* is also expressed in PT tissue in a genetic background-dependent manner in the PyMT-F1 cohort (Fig S8d). Based on these data, we conclude that the MSGE of these models is largely oncogenic driver-dependent.

Similarly, GO analysis revealed that there were no Lung met-specific gene expression pathways common across all oncogenic driver models (Fig S8e and Table S12). However, it was noted that the Myc model MSGE only enriched a single pathway, the *Wnt Signaling pathway*. Three pathways, *Complement and coagulation cascades, Platelet activation,* and *Malaria,* were significantly enriched and shared by the Her2, C3Tag, and PyMT models (Fig 5k). Interestingly, while these pathways are related by their involvement with red blood cell biology, the gene lists that define them overlap less than 7% (Fig S8f). Upon assessment of the genes within each pathway, we noted that Her2, C3Tag, and PyMT enrich largely independent gene sets except for the cell adhesion molecules *Icam1, Vwf, and Itgb2* (Fig 5l and Table S13).

RNA-seq data was next analyzed for metastasis-specific changes in transcript levels to assess model–specific splicing. Upon comparison of Lung mets and PTs from the oncogenic driver cohort, we identified a single transcript significantly altered in metastases compared to PTs and common to all models (Fig S9a and Table S14). *Rnase4* isoform 2, and not isoform 1, was significantly increased in metastatic tissue compared to PT (Fig S9b-c). *Rnase4* isoform 1 and 2 share the same promoter and coding exons but have unique 5’ untranslated regions (UTRs)^22,23^. We noted that overall expression of *Rnase4* remained unchanged, which may be explained by a compensatory, although non-significant, decrease of isoform 1 (Fig S9b). We asked if Rnase4 might also be alternatively spliced in lung metastases across the PyMT-F1 cohort. A statistically significant increase in levels of isoform 2 was observed in BALB, SEAGN, and BL6, but not any of the remaining strains (Fig S9d). Comparative analysis of splicing across the PyMT-F1 cohort revealed no commonly altered transcripts, suggesting that metastasis-specific splicing may be determined in-part by genetic background (Fig S9e).

We next asked how genetic background could shape gene expression programs in macro-metastases. Upon visualization of PT and Lung met gene expression by unsupervised PCA for each PyMT-F1 strain, we observed no distinct separation between tissue types for BL6, MOLF, and CAST (Fig 5m-o), but distinct clustering of samples by tissue in BALB, SEAGN, and BL10 (Fig 5p-r). The heterozygous PyMT F1s had a higher degree of MSGE compared to homozygous FVB PyMT samples, with the number of altered genes ranging between 814 and 3226 for BL6, BL10, SEAGn, MOLF, and CAST (Fig 5s-t and Table S11). Comparison of the MSGE between the F1s using a six-part Venn diagram analysis showed that close genetic background did not always correspond with highest overlap in MSGE (Fig S10a). MOLF, BL6, and BALB MSGE were most similar to their corresponding strain pairs, CAST, BL10, and SEAGN, respectively, with about 50% of the MSGE in these strains overlapping with the MSGE of their closely related counterparts. However, SEAGN had the highest overlap in MSGE with CAST, while CAST and BL10 strains had a higher percentage of overlap with one another. Furthermore, we noted that MSGE was also highly individualized between F1 strains, with unique changes in gene expression accounting for 23-58% of MSGE (Fig S10a). Only 60 genes were common to MSGE of all strains (1.9-13.6% of total MSGE for each F1 strain), and these were enriched for extracellular matrix, cell adhesion, autophagy factors, and glycoproteins as determined by DAVID analysis (Table S15)^24,25^.

Further GO analysis was performed using the MSGE for each PyMT-F1 strain (Table S16). Venn diagram analysis of the enriched pathways revealed *ECM-receptor interaction*, *Gap junction*, *Focal adhesion*, and *Renin Secretion* as four pathways commonly enriched by MSGE across all PyMT-F1s (Fig 5u). The genes included in the KEGG *ECM-receptor interaction* and *Gap junction* pathways overlap with the *Focal adhesion* pathway by 74% and 30%, respectively, but not one another (Fig S10b). As such, when we compared the MSGE of each pathway between strains we identified four commonly altered genes (*Thbs2, Col1a2, Vwf,* and *Egflam*) from the *Focal Adhesion* pathway that were also found within either the *ECM receptor interaction* or *Gap junction* pathways (Fig 5v and Table S17). Furthermore, three genes (*Adrb2, Npr1,* and *Acer2*) from the *Renin Secretion* pathway were upregulated in the MSGE of all F1 strains (Fig 5w). Based on GTex project data, none of these commonly altered genes are Lung-specific, all are expressed in breast tissue, and some are ubiquitously across many tissues (Fig S10c). In summary, while the same pathways were enriched by all strains, each F1 appears to modulate gene expression differently to achieve enrichment of these pathways.

### MSCNV and MSGE from both cohorts target common cellular processes

Analyses conducted in this study sought to identify commonalities between models. However, in many instances, particularly among PyMT-F1s, the models were more unique than similar. Indeed, we noted that the oncogenic driver models had 38-72% of MSCNV enriched pathways that were unique to each model, and the altered genetic background of the PyMT-F1 cohort resulted in no commonly enriched genes or pathways from MSCNV, but rather 60-88% unique pathways in each F1 strain (Fig 2a and Fig 3a). To visualize and summarize the differences between models the top 70 GO terms from MSCNV gain and loss were sorted into thirteen categories and the percent of each category within the top pathways presented as heatmaps (Fig 6a and Table S7 and S9). Immune (cyan) and metabolic (pink) pathways were dominant across all four oncogenic driver models, in both MSCNV gain and loss. However, each model displayed a unique distribution of pathways, and MSCNV gain and loss were also distinct within the same model, which was most pronounced in the C3Tag model (Fig 6a). MSCNV loss among the PyMT-F1 BALB, AKR, and SEAGN models most closely resembled the distribution observed in the oncogenic driver models, with immune and metabolic pathways dominant, but developmental pathways (grey) were the most significantly enriched across this cohort. From this analysis, we conclude that despite exploiting the same oncogenic driver, genetic background has a stronger influence on the genes enriched within MSCNV (Fig 6a).

**Figure 6.**
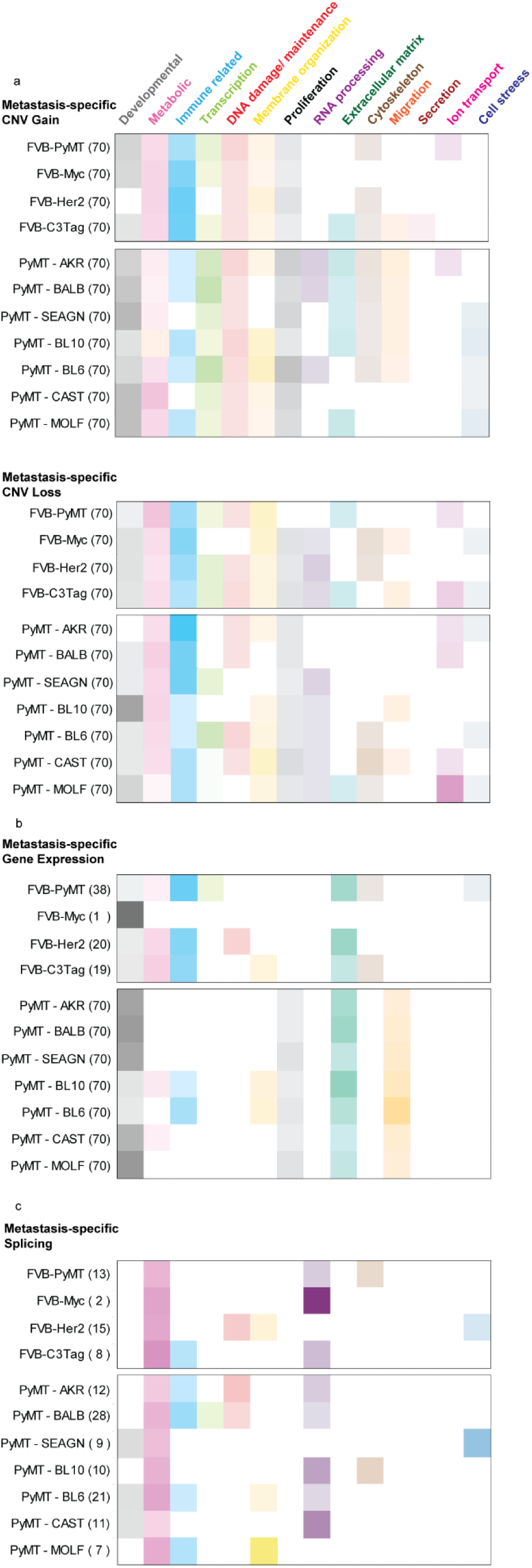
Oncogenic driver and genetic background shape unique MSCNV and MSGE to target common cellular processes. (a-c) Categorization of the top 70 (or otherwise indicated) significantly enriched gene ontologies from MSCNV (a), MSGE (b), and lung met-specific alternative splicing (c). Uncategorized ontologies were excluded from the heat maps. Color intensity represents the percent of the analyzed pathways that fell within that category.

We next extended this analysis to visualize differences in MSGE from RNA-seq data (Fig 6b). We again observed an enrichment of immune pathways across the oncogenic driver models, except for in the Myc model, which had limited MSGE and only a single enriched pathway (Fig S8a and Table S12 and S16). For the PyMT-F1 cohort, we observed more similarity in metastasis-specific gene expression pathways across strains, compared to the wide variety enriched by CNV (Fig 6a-b). We noted that despite seeing no common developmental pathways, this category made up the majority of MSGE-enriched ontologies among five of the models. In addition, ECM, migration, and proliferative pathways were enriched to varying degrees in the MSGE of each PyMT-F1 strain. Interestingly, the MSGE in the PyMT-F1 cohort represented fewer categories compared to the MSCNV-enriched pathways. Overall, due to the low number of genes and pathways common to MSGE across all PyMT-F1 strains, we conclude that while genetic background can have a profound impact on gene expression within metastatic lesions, these changes may ultimately impact the same cellular processes. This analysis demonstrated that a more diverse genetic background, modeled by the heterozygous PyMT-F1 cohort, shifted MSGE to more developmental and less immune-related ontologies. However, Extracellular matrix (green) genes appear to be commonly modulated by MSGE across all models.

Finally, we assessed pathways enriched by metastasis-specific splicing. In both the oncogenic driver and PyMT-F1 cohorts, we noted that fewer pathways were significantly enriched by alternatively spliced genes compared to MSCNV and MSGE (Fig 6c and Table S18). Despite there being no common alternatively spliced transcripts across the eleven models used in this study, we noted that metabolic and RNA processing (purple) ontologies were dominant in all models. However, each model had a unique distribution of pathways significantly enriched by metastasis-specific splicing programs.

## Discussion

Metastasis is a complex phenotype, driven by the interplay of tumor cell intrinsic, microenvironmental, and systemic factors^26^. Classic cytogenetics studies and more recent sequencing-based work has shown that metastases usually arise from a single, or at most a limited number, of subclones within the primary tumor^27^. In addition, clinical and experimental data suggest that only a small fraction of disseminating tumor cells are able to form secondary lesions^28^. These and other data have been interpreted to suggest that tumors evolve over time until a subset of tumor cells acquire all of the necessary attributes to successfully complete the metastatic cascade.

To date, however, in contrast to what has been observed in primary tumors, sequencing studies in clinical samples have not revealed the presence of high frequency, reproducible SNVs associated with metastatic progression^29–31^. In contrast, previous studies from our laboratory, comparing matched primary and metastatic lesions from the highly metastatic MMTV-PyMT mouse model of breast cancer, did identify reproducible metastasis-associated point mutations in *Shc1* and *Kras*^6^. These studies, however, were performed on a much more genetically homogeneous set of animals than is observed in the human population, suggesting that these potential metastasis driver mutations might only represent a small fraction of the patient population. In addition, these observations were made for a single oncogenic driver model, PyMT, rather than the multiple oncogenic initiating events observed for human patients.

To better understand how genomic evolution may contribute to metastatic progression in commonly used GEMMs and to compare them to the expanding clinical data, we have extended our previous work to include two additional models of breast cancer metastasis, the C3(1)TAg and MMTV-Myc models, and expanded on the previously analyzed MMTV-Her2 model. Moreover, for the MMTV-PyMT model, we have expanded the repertoire to include pairs of phylogenetically related strains with significantly different inherited propensity to metastasize to include population diversity into the analysis. The total data set analyzed in this study consists of 197 matched primary and metastatic lesions spanning models of luminal, Her2, and basal-like breast cancer.

Consistent with our previous study, analysis of the expanded data set failed to reveal high frequency SNVs associated with metastasis across all of the models. However, a total of 13 genes were identified that were mutated in 2 or more independent animals, The most frequently observed gene was *Kras*, which had metastasis-associated activating mutations in 5 out of 197 animals. All of the remaining genes with metastasis-associated SNVs were present in only 2 or 3 animals. Reproducible metastasis-associated SNVs were not observed in 84% (166/197) of the animals. This suggests that either SNVs do not play a significant role in metastatic progression or that there may be many SNVs that contribute at a low frequency to metastatic progression. The ability to identify the low frequency metastasis-associated SNVs in the mouse model, compared to human clinical samples, may be due to species-specific effects, which cannot be excluded at this time. The mouse data generated here, however, is consistent with current clinical data suggesting that there is a lack of activating mutations for precision medicine for anti-metastatic therapies that would be of benefit for significant fractions of breast cancer patients.

We subsequently performed CNV analysis on the data to determine whether there were common CNVs that might contribute to metastatic progression. Analysis revealed that oncogenic drivers on the FVB homozygous genetic background had similar degrees of CNV in both primary and secondary lesions, though there were differences in the size of the CNVs between models. Further, in both PTs and lung mets we observed some strains with high numbers of small events and others with small numbers of larger events, accounting for the similar degree of copy number change overall. Notably, the size of copy number events in this study are within the range observed across cancer types by Steele et al. in a recently published analysis of copy number signatures from clinical data^32^. The major determinant of CNV variation, however, was genetic background. This was most apparent in the comparison of the phylogenetically related BALB and SEAGn strains. The SEAGn strain was generated by a cross between BALB and PJ mice, followed by 20 generations of brother-sister mating. This congenic strain genome is 50% BALB yet shows significantly less genomic stability compared to the BALB primary tumor. Together this data suggests that genetic background may direct the mechanisms leading to copy number events.

Further analysis focused on metastasis-associated events revealed common CNVs, associated genes, and gene ontologies across the oncogene driver models on the FVB homozygous background. However, these common metastasis-associated features are lost when the genetic background was varied in the MMTV-PyMT cohort. This suggests that while the analysis of the four homozygous FVB oncogenic models may be valuable for identification of some common metastatic mechanisms, they are unlikely to represent the majority of the human patient population. Moreover, consistent with our previous observations of inherited metastatic susceptibility, this suggests that an individual’s personal ancestry is a major factor in metastatic genome evolution. In addition, these data suggest that there are likely multiple paths that tumor cells utilize to achieve metastatic competency, which indicates that multiple strategies to prevent or treat established lesions may be required across the human population.

Examination of the data at a broader level provides cause for optimism. Although individual Gene Ontology terms from CNV and transcriptome analysis were not shared across the models and genotypes, grouping individual ontologies into broader categories demonstrates potential commonalities that might be targeted. When considered as broad categories, developmental pathways, ECM, RNA biology, and metabolism appear to be common metastatic categories across all of the animals, although different Gene Ontologies are affected in each of the cohorts. Each of these broad categories have already been previously associated with metastatic disease. The data presented here suggests that broader investigations into alternative paths or ontologies within each of these categories may be necessary to achieve higher efficacy of anti-metastatic therapies.

In summary, here we have demonstrated that 1) Genetic background plays a role in primary tumor stability and the size of copy number event regions, 2) Genetic background is a strong determinant of which genes are specifically enriched in CNV. 3) Oncogenic driver and genetic background shapes gene expression in metastatic tumors, and 4) Metastasis-specific ontology clusters, rather than individual Gene Ontologies, identify common biological programs in metastatic disease that might be considered for clinical intervention. An increased understanding of the complex interplay between these factors will provide guidance for the improved use and interpretation of these models in relation to human patient populations for the investigation of progressive metastatic disease.

## Methods

### Ethics statement

The research described in this study was performed under the Animal Study Protocol LPG-002, approved by the National Cancer Institute (NCI) Animal Use and Care Committee. Animal euthanasia was performed by cervical dislocation after anesthesia by Avertin.

### Genetically engineered mouse models and tissue isolation

FVB/N-Tg(MMTV-PyVT)634Mul/J (PyMT), FVB/N-Tg(MMTVneu)202Mul/J (Her2) male mice were obtained from The Jackson Laboratory. FVB/N-Tg(C3(1)-TAg) (C3-TAG) and FVB/N-Tg(MMTV-Myc) (MYC) male mice were a generous gift from Dr. Jeffrey Green (NCI, Bethesda, MD). Male PyMT mice were crossed with female wild type FVB/NJ, MOLF/EiJ, CAST/EiJ, C57BL/6J, C57BL10/J, BALB/cJ, Sea/GnJ, and AKR/J mice, which were also obtained from The Jackson Laboratory. Male Her2, C3-TAG, and MYC mice were crossed with female wild type FVB/NJ mice.

All female F1 progeny were genotyped by the Laboratory of Cancer Biology and Genetics genotype core for the PyMT, Her2, Myc, or C3-TAG gene and grown until humane endpoint. Mice were euthanized using intraperitoneal Avertin to anesthetize followed by cervical dislocation. All primary tumors generated by one animal were isolated, weighed, randomly sampled, and combined into a single cryovial. Metastatic nodules and normal (tail) tissue were also isolated immediately following euthanasia, placed in cryovials, and snap frozen in liquid nitrogen. Tissue samples were then stored at -80°C.

### gDNA and RNA isolation

The combined primary tumor tissue from one mouse was ground on dry ice and small fragments were taken for nucleotide extraction isolation. Whole metastases were used for gDNA or RNA isolation.

For gDNA isolation, tissue was lysed using Tail Lysis Buffer (100 mM Tris-HCl pH 8.0, 5 mM EDTA, 0.2% SDS, 200 mM NaCl, 0.4 mg/ml proteinase K) at 55°C overnight. Samples were then placed in a shaking (1400 rpm) heat block for 1 hour at 55°C. RNaseA (Thermo Fisher Scientific) was added (2 mg/ml final) and lysates were incubated on the bench for 2 minutes. gDNA was then isolated using the ZR-Duet DNA/RNA MiniPrep kit (Zymo Research).

For RNA isolation, the tissue was mechanically dissociated using a tissue grinder while submerged in 1 ml of TriPure (Roche). 200 μl of chloroform (Sigma-Aldrich) was then added and the soluble fraction was isolated by centrifugation at 12,000 rpm for 15 minutes at 4°C. RNA was then precipitated with the addition of 500 μl isopropanol and incubation of the sample at −20°C for 2 hours. Pure RNA was then extracted using the RNA: DNA mini-prep kit (Zymogen) and samples were eluted in 100 μl (PT) or 50 μl (metastases) of DEPC water (Quality Biological). RNA was isolated from cell lines using TriPure as described above, but, following isopropanol precipitation, RNA was washed with 75% ethanol (Sigma-Aldrich) followed by 95% ethanol before being resuspended in 100 μl of DEPC water.

### Sequencing and analysis

All analyses were carried out on the NIH Biowulf2 high performance computing environment. All analyses were performed using software default parameters if not otherwise specified.

#### Exome sequencing and data analysis

Exome sequencing was performed by the NCI Center for Cancer Research (CCR) Genomics Core and the NCI Illumina Sequencing Core. Exome libraries were prepared by the Genomics Lab using Agilent SureSelectXT Mouse All Exon target enrichment kit. Libraries were barcoded and pooled before sequencing on an Illumina HiSeq3000 or HiSeq4000 to an average depth of 40x. Samples were trimmed of adapters using Trimmomatic software. Bam files were uploaded into Nexus Copy Number 10.0 (BioDiscovery) and processed using SNP-FASST2 Segmentation. The trimmed reads from tumor gDNA were aligned to normal tail gDNA sequence from each model, and then to the mouse NCBI build 38 (mm10) reference genome.

#### Single nucleotide variant analysis

This analysis was performed as described in our previously published work^6^*. Copy number analysis*

Following initial data processing, the *Total copy number events* and *% of the genome changed* by copy number for each sample were downloaded from the Data Set tab. Aggregate data from PTs and lung mets was attained by selecting only the sample from the desired tissue type within the Data set tab, and then selecting *View* and then *Aggregate.* Aggregate regions of CNV were defined as having a p value cutoff below 0.05 and an aggregate percent frequency of 35% and above. Region length, event type, number of genes, and gene names were exported as .txt from aggregate analysis. Average region length, total number of events and total number of genes were calculated using Excel (Microsoft).

For lung met-specific CNV analysis, the *Classic* comparison tool was utilized in Nexus. In the Data Set tab, all samples from one mouse model were selected before navigation to the *Comparisons* tab. From here, a comparison was added using *Tissue* as the distinguishing factor, and *Average* was selected as the comparison baseline. Upon selection of the comparison and then *View,* lung met specific *e*vent type, region length, p value, number of genes, and gene names were exported as .txt files. Average region length, total number of events, and total number of genes were calculated using Excel. By highlighting specific CNV events within the comparison V*iew* tab, a pathway enrichment analysis was performed for the genes found within regions of CN Gain or CN Loss. A list of enriched *Biological Processes* was exported as a .txt file including p values. The percent CN gain, loss, or allelic imbalance in the primary tumors and lung mets for each gene within each pathway was obtained by selecting the individual biological process name. Calculation of % Lung met gain or loss - % PT gain or loss was performed in Excel and heat maps were created using the Excel conditional formatting tool.

#### RNA sequencing and data analysis

RNA quality was tested using the Agilent 2200 TapeStation electrophoresis system, and samples with an RNA integrity number (RIN) score >7 were sent to the Sequencing Facility at Frederick National Laboratory. Preparation of mRNA libraries and mRNA sequencing was performed by the Sequencing Facility using the HiSeq2500 instrument with Illumina TruSeq v4 chemistry.

RNA-seq data was analyzed using Partek Flow software (Kanehisa Laboratories). RNA-seq reads were uploaded into Partek Flow, aligned with the mouse mm9 genome assembly, gene counts were then determined, and then normalized. Principal Component Analysis was performed using unsupervised principal component analysis for either all samples, or samples filtered by mouse model. Differentially expressed gene lists were generated Gene Specific Analysis (GSA) for FVB-Driver models and Dseq2 for PyMT-F1 models. Differential gene expression was defined as a fold change with a false discovery rate less than 0.05. The Differential expression of genes within regions of CNV was defined as fold change with FDR less than 0.1.Volcano plots for Lung met v PT gene expression were created using unfiltered GSA results for all models. Gene ontology analysis was performed using differentially expressed gene lists and the Partek Flow Gene Set Enrichment tool. Pathways enriched with a p value less than 0.05 were considered significant. Gene ontology categories were assigned manually to the top 70 pathways or if less than 70 pathways were significantly enriched then all pathways with p<0.05 were included. Heatmaps were created using the Excel conditional formatting tool, where color intensities were determined by the percent of the enriched terms found within each category out of the total terms categorized.

Differential splicing analysis was performed using Partek flow. Following alignment to mm9 genome assembly, transcript counts were normalized and then each model was analyzed using the Partek tool to detect all splicing and differential splicing in Lung mets v PT. Alternative splicing was defined as both transcript differential expression with p value less than 0.05 and an alt-splicing p value less than 0.05. Using these p value cuts-offs, alternatively spliced gene lists were analyzed by Partek Gene Set Enrichment to identify pathways enriched by lung met alternatively spliced genes. Significantly enriched pathways were defined as those with p values less than 0.05.

## Supplemental Tables

Table S1 - Sample numbers and attributes

Table S2 - Total number of CN aberrations and % genome with CNV for all models.

Table S3 - Type, size, and loci for aggregate CNVs in FVB-driver model PTs (Sheet 1) and Lung mets (Sheet 2), and for PyMT-F1 cohort PTs (Sheet 3) and Lung mets (Sheet 4).

Table S4 - Type and length of MSCNV for FVB-driver (Sheet 1) and PyMT-F1 (Sheet 2) cohorts.

Table S5 - MSCNV gene lists for FVB-driver (Sheet 1) and PyMT-F1(Sheet 2) cohorts.

Table S6 - DAVID functional clustering annotation analysis using 1676 common MSCNV gene list from FVB driver models.

Table S7 - GO analysis for genes within MSCNV for each FVB driver model. PyMT (Sheet1), Her2 (Sheet 2), C3Tag (Sheet 3), Myc (Sheet4). Overlapping GO terms with assigned category and -Log10(p values) (Sheet 5).

Table S8 - FVB Driver Model Heat Maps separated by pathway category. Immune Pathways (Sheet1), Metabolic Pathways (Sheet2), Developmental pathways (Sheet 3), and Other Pathways (Sheet4). Blue indicates genomic gain and red indicates genomic loss in metastatic tissue vs PT.

Table S9 - GO analysis for MSCNV-enriched genes for each F1 of the PyMT-F1 cohort. Separated by sheets 1-7.

Table S10 - Gene expression overlaps with MSCNV for each model.

Table S11 - MSGE for each individual model separated by sheets 1-11.

Table S12 - Oncogenic Driver cohort GO enrichment in MSGE for each driver model separated by sheets 1-4.

Table S13 - MSGE of FVB-PyMT, Her2, and C3Tag common pathways as heatmaps. Indicating upregulation (green), down regulation (yellow), and significance (red text) for each gene within each model.

Table S14 - Metastasis-specific splicing for each individual model separated by sheets 1-11.

Table S15 - DAVID functional clustering annotation analysis using PyMT-F1 MSGE overlap.

Table S16 - PyMT-F1 cohort GO enrichment in MSGE for each F1 model separated by sheets 1-6

Table S17 - MSGE of F1-PyMT common pathways as heatmaps. Indicating upregulation (green), down regulation (yellow), and significance (red text) for each gene within each model.

Table S18 - GO analysis of MS splicing for each model separated by sheets 1-11.

Table S19 – Statistical summary of data analyses in Figure 1 and Figure S3.

