## Supplemental figures for "Investigating the genomic landscape of mouse models of breast cancer metastasis"

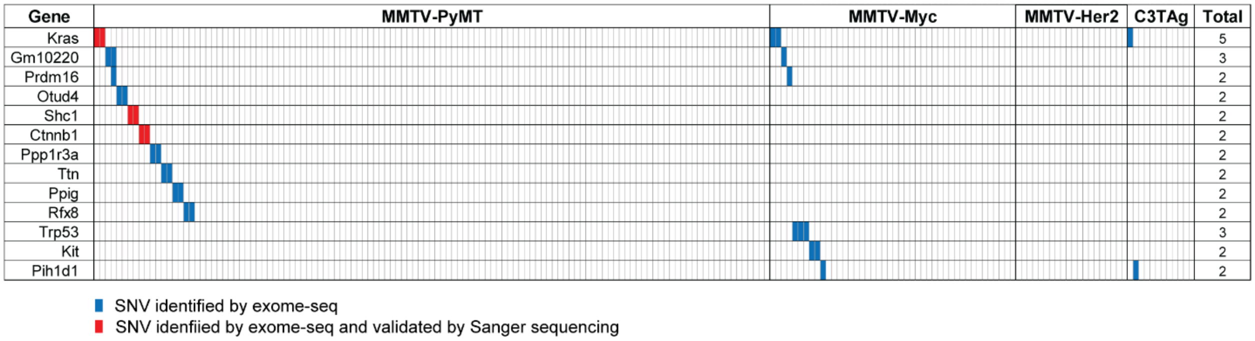


Supplementary Figure 1 Metastasis-specific SNV oncoprint summary

SNV analysis summary across oncogenic driver and PyMT-F1 cohorts. Each box represents a mouse, with oncogenic driver indicated across the top adjacent bars. Genes with metastasis-specific high allele frequency SNVs are listed on the left, and colored boxes indicate a mouse having a metastasis-specific SNV with allele frequency above 0.6. Red identifies SNVs identified by exome-seq, and blue indicates an additional level of validation with Sanger sequencing. The total number of mice with a metastasis-specific SNV for each gene is listed on the right.

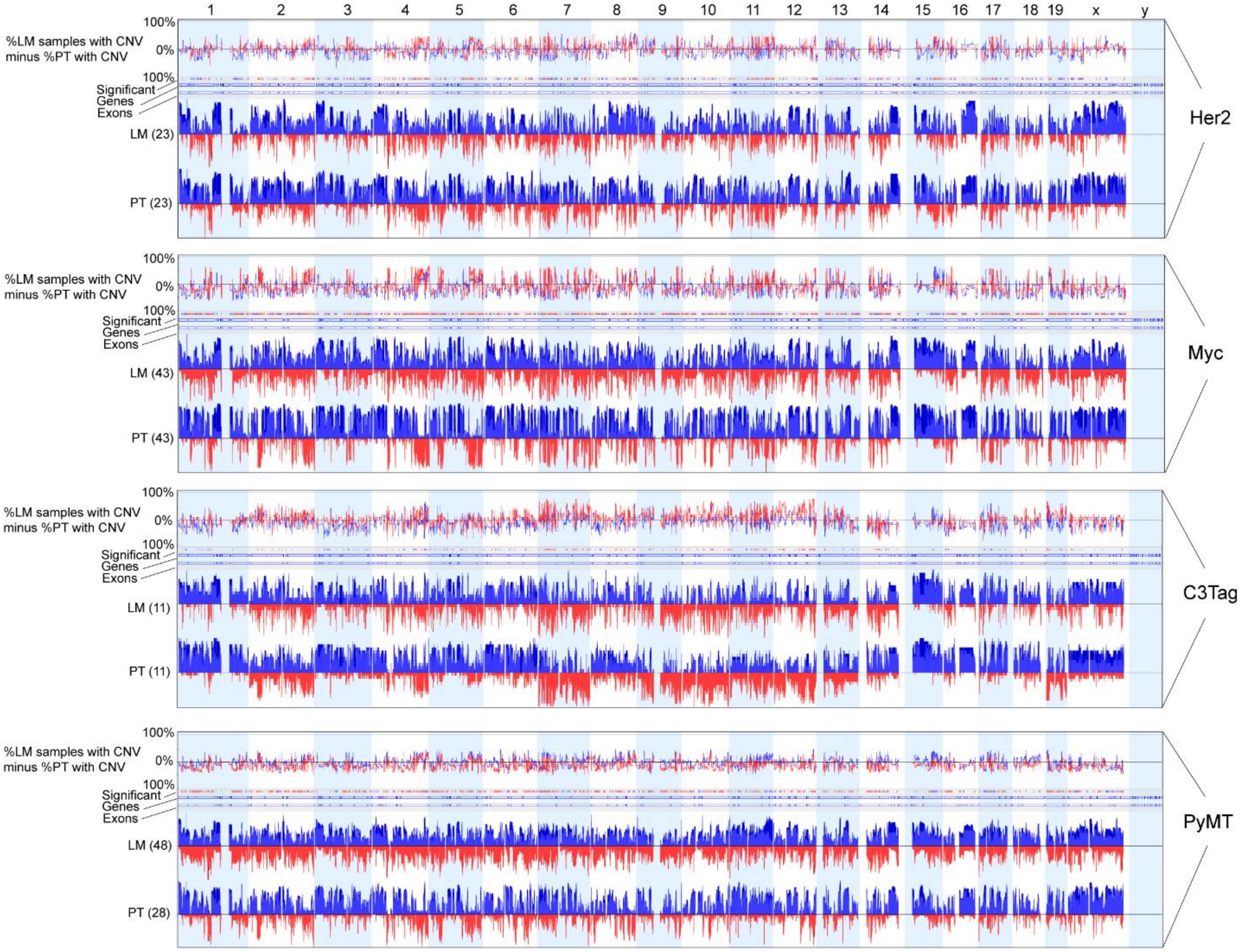


Supplemental Figure 2 FVB driver model PT and Lung met CNV

Whole genome aggregate CNV traces for PTs and lung mets from each model, also showing a trace of % CNV gain or loss in lung mets minus % CNV gain or loss in PT, and regions with significant differences ($p<0.05$). Blue indicates CNV gain and red indicates CNV loss. The number of tissue samples for each model is listed with “LM” and “PT”.

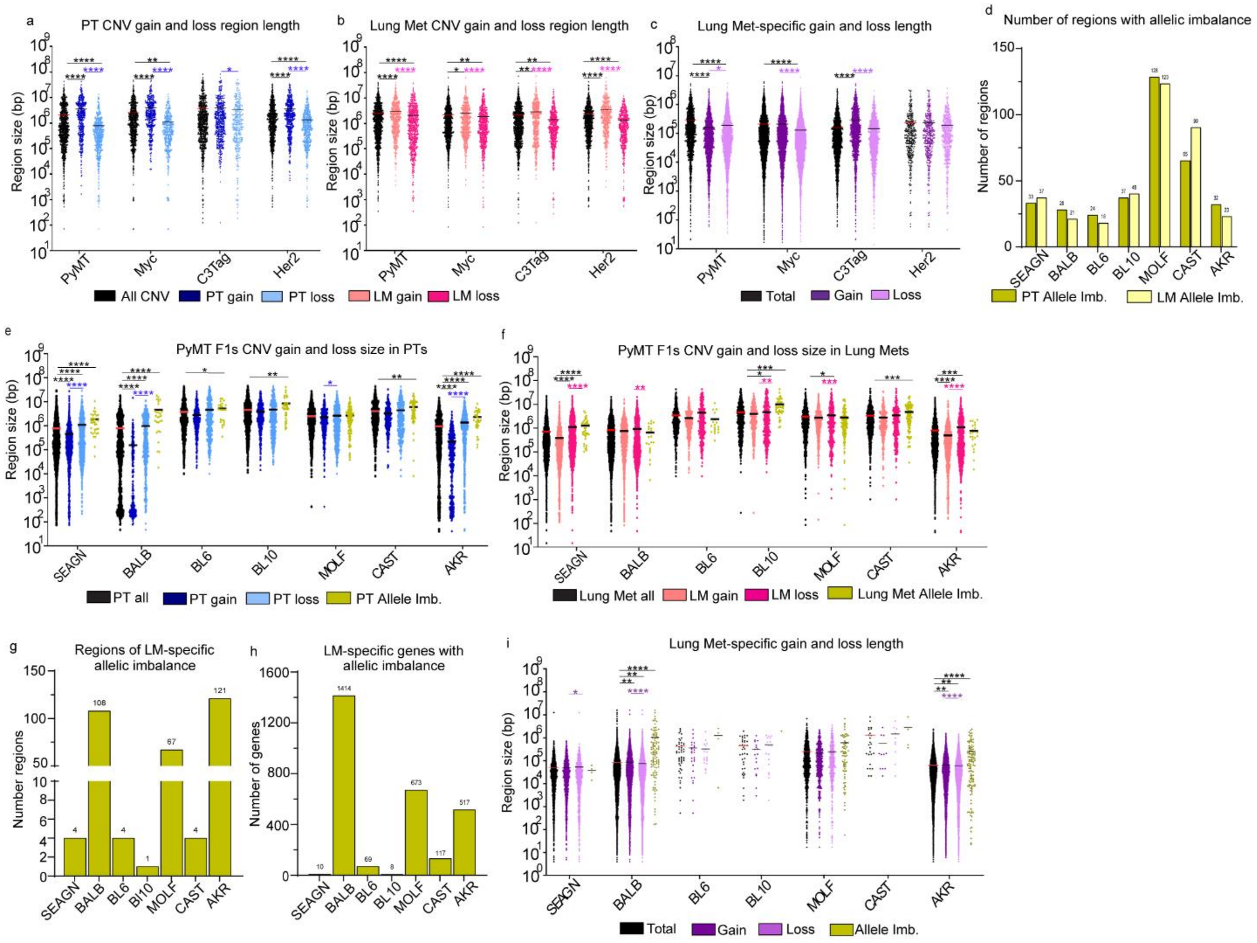


Supplemental Figure 3 CNV region size by model, tissue, and CNV type

(a-c) Dot plots showing length of individual regions of CNV separated by FVB driver model, showing all CNV (black), with PT CNV gain (dark blue) and PT CNV loss (light blue) shown in (a), Lung met (LM) CNV gain (light pink) and LM CNV loss (dark pink) shown in (b), and MSCNV gain (dark purple) and loss (lilac) shown in (c). (d) Histogram showing number of regions of allelic imbalance in the PyMT-F1 model PT (gold) and LM (yellow). (e-f) Dot plots showing length of individual regions of CNV separated by PyMT-F1 model, showing all CNV (black), with PT CNV gain (dark blue) and PT CNV loss (light blue) shown in (e) and LM CNV gain (light pink) and LM CNV loss (dark pink) shown in (f). (g-h) Histograms showing the number of regions of allelic imbalance due to MSCNV (g) and the number of genes found within those regions (h). (i) Dot plot showing MSCNV region sizes in total (black) and separated according to regions of gain (dark purple), loss (lilac), and allelic imbalance (gold). Significance determined by Kruskal-Wallis test, *(p<0.05), **(p<0.01), ***(p<0.001), ****(p<0.0001).

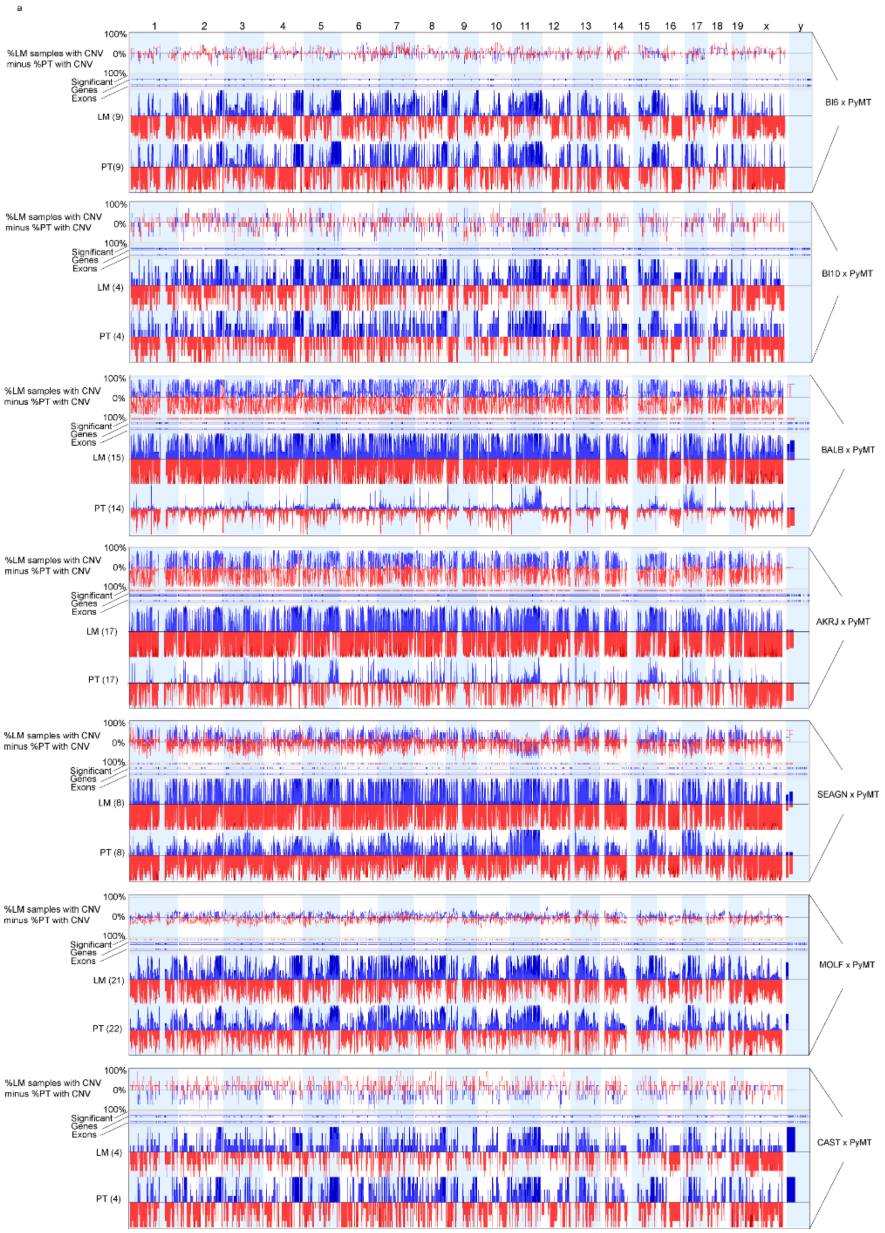


Supplemental Figure 4 PyMT-F1 model PT and Lung met CNV

(a) Whole genome aggregate CNV traces for PTs and lung mets from each model, also showing a trace of % CNV gain or loss in lung mets minus % CNV gain or loss in PT, and regions with significant differences ($p<0.05$). Blue indicates CNV gain and red indicates CNV loss. The number of tissue samples for each model is listed with “LM” and “PT”.

a

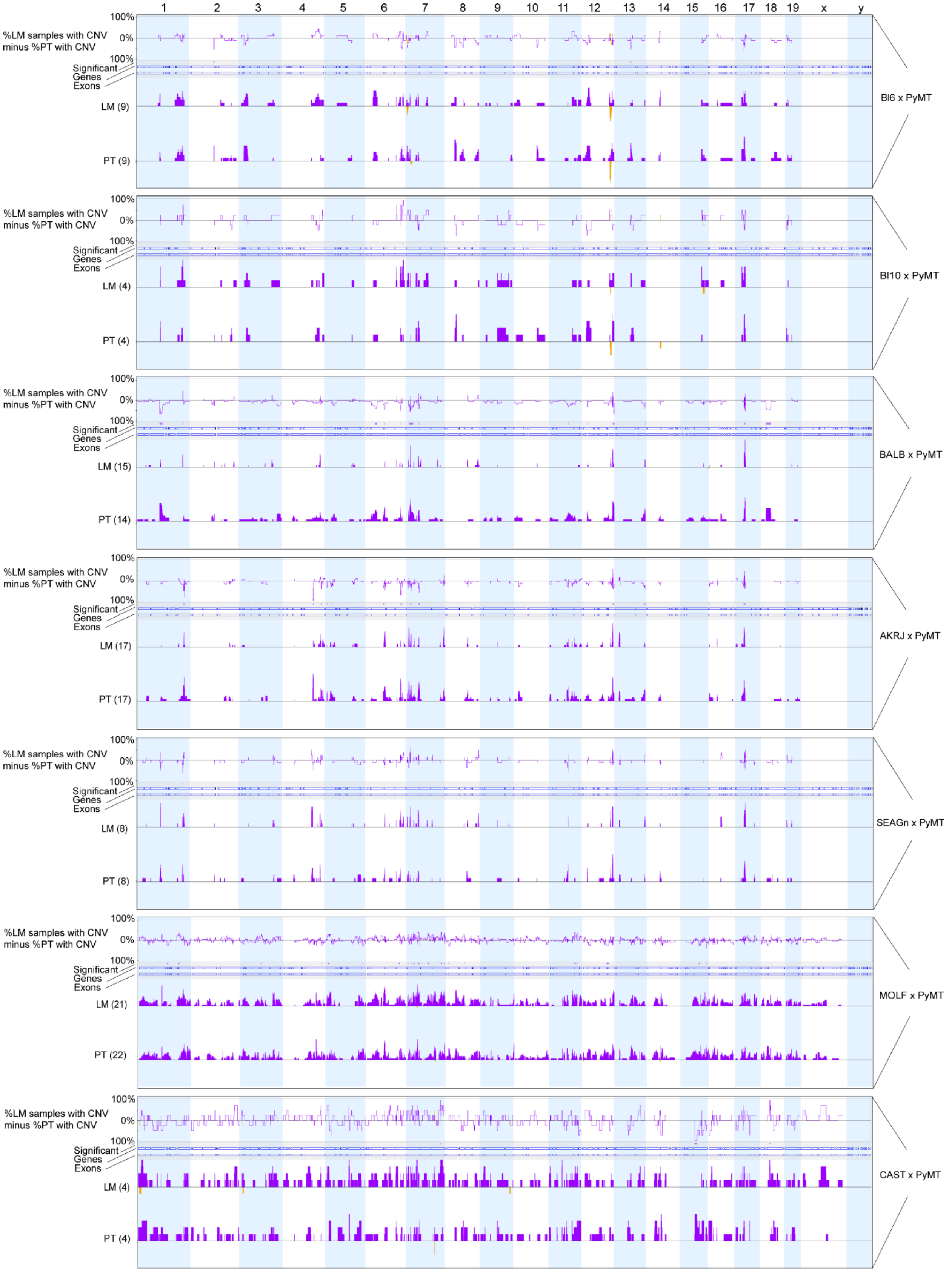
1 2 3 4 5 6 7 8 9 10 11 12 13 14 15 16 17 18 19 x y
%LM samples with CNV minus %PT with CNV
100%
0%
100%
Significant
Genes
Exons
LM (9)
PT (9)
Bl6 x PyMT
LM (4)
PT (4)
Bl10 x PyMT
LM (15)
PT (14)
BALB x PyMT
LM (17)
PT (17)
AKRJ x PyMT
LM (8)
PT (8)
SEAGn x PyMT
LM (21)
PT (22)
MOLF x PyMT
LM (4)
PT (4)
CAST x PyMT

Supplemental Figure 5 PyMT-F1 model PT and Lung met allelic imbalance

(a) Whole genome aggregate CNV traces for PTs and lung mets from each model, also showing a trace of % allelic imbalance in lung mets minus % CNV allelic imbalance in PT, and regions with significant differences (p<0.05). The number of tissue samples for each model is listed with “LM” and “PT”.

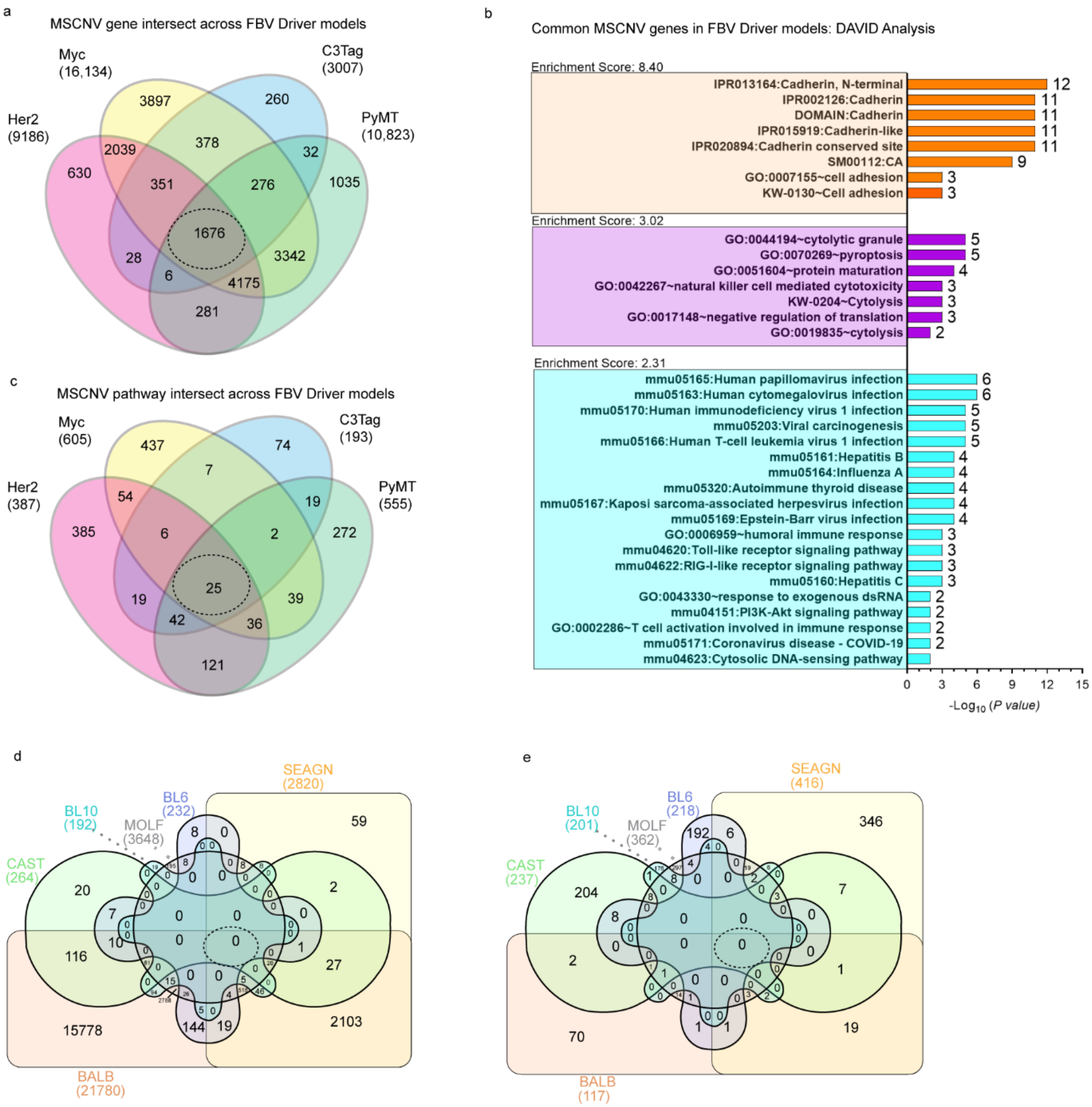


Supplemental Figure 6 CNV-impacted genes and ontologies across models

(a) Venn diagram analysis of MSCNV-altered genes from the FVB driver models with overlap indicated by the dotted-line circle. (b) DAVID analysis summary of overlapping genes indicated by the dotted-line circle in (a). (c) Venn diagram of the pathway analysis for MSCNV in each individual FVB driver model, with overlap indicated by the dotted-line circle. (d-e) Venn diagram analysis of MSCNV-altered genes (d) and ontologies (e) from the PyMT-F1 models with overlaps indicated by the dotted-line circles.

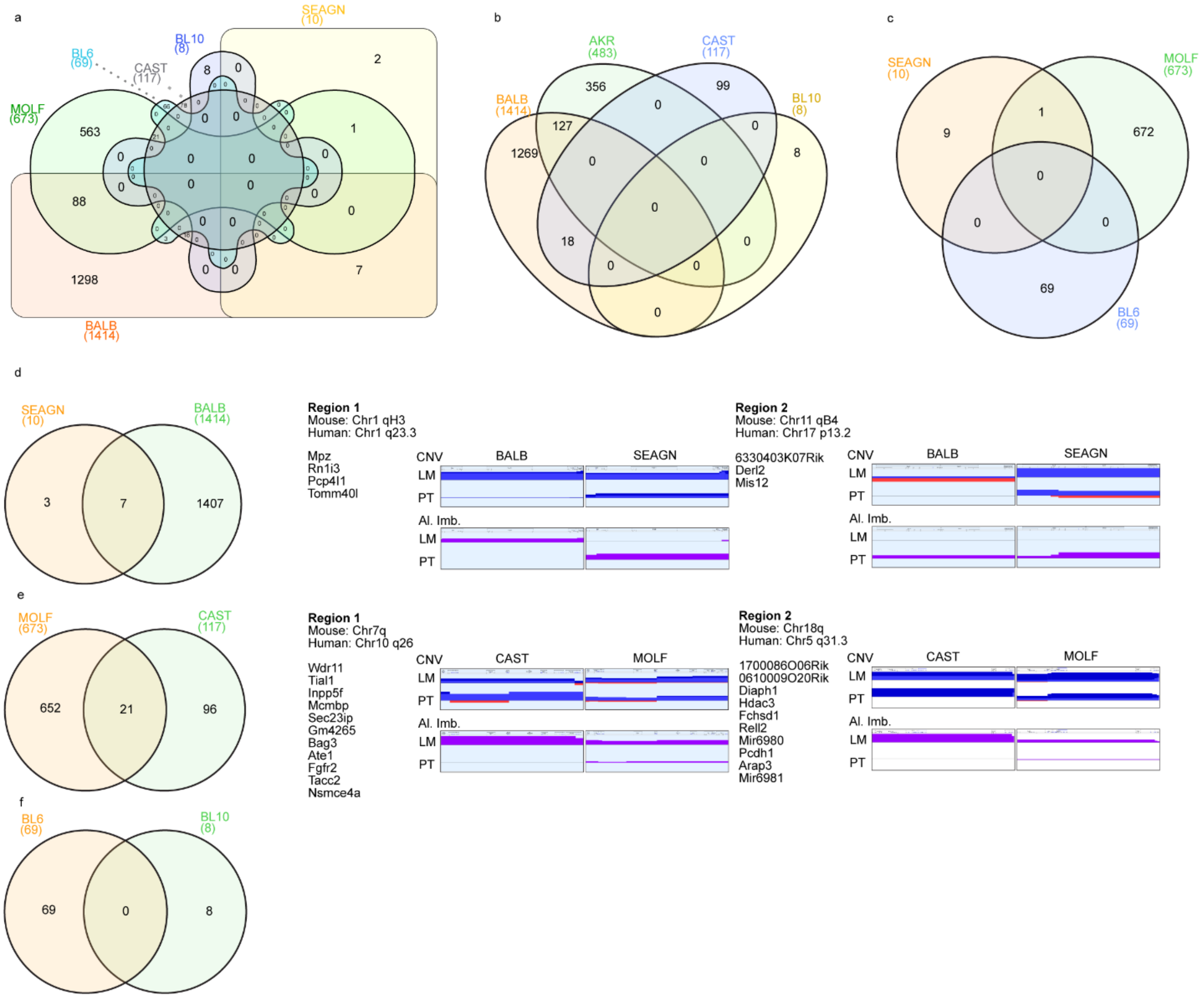


Supplemental Figure 7 Common allelic imbalance between F1-PyMT model

(a) Venn diagram analysis of gene lists within metastasis-specific allelic imbalance for each model. (b-c) Venn diagram analysis of gene lists within metastasis-specific allelic imbalance for high metastatic efficiency models (b) and low metastatic efficiency models (c). (d) Venn diagram analysis of SEAGN and BALB metastasis-specific allelic imbalance with PT and LM CNV traces from common regions of imbalance. (e) Venn diagram analysis of MOLF and CAST metastasis-specific allelic imbalance with PT and LM CNV traces from common regions of imbalance. (f) Venn diagram analysis of BL6 and BL10 metastasis-specific allelic imbalance.

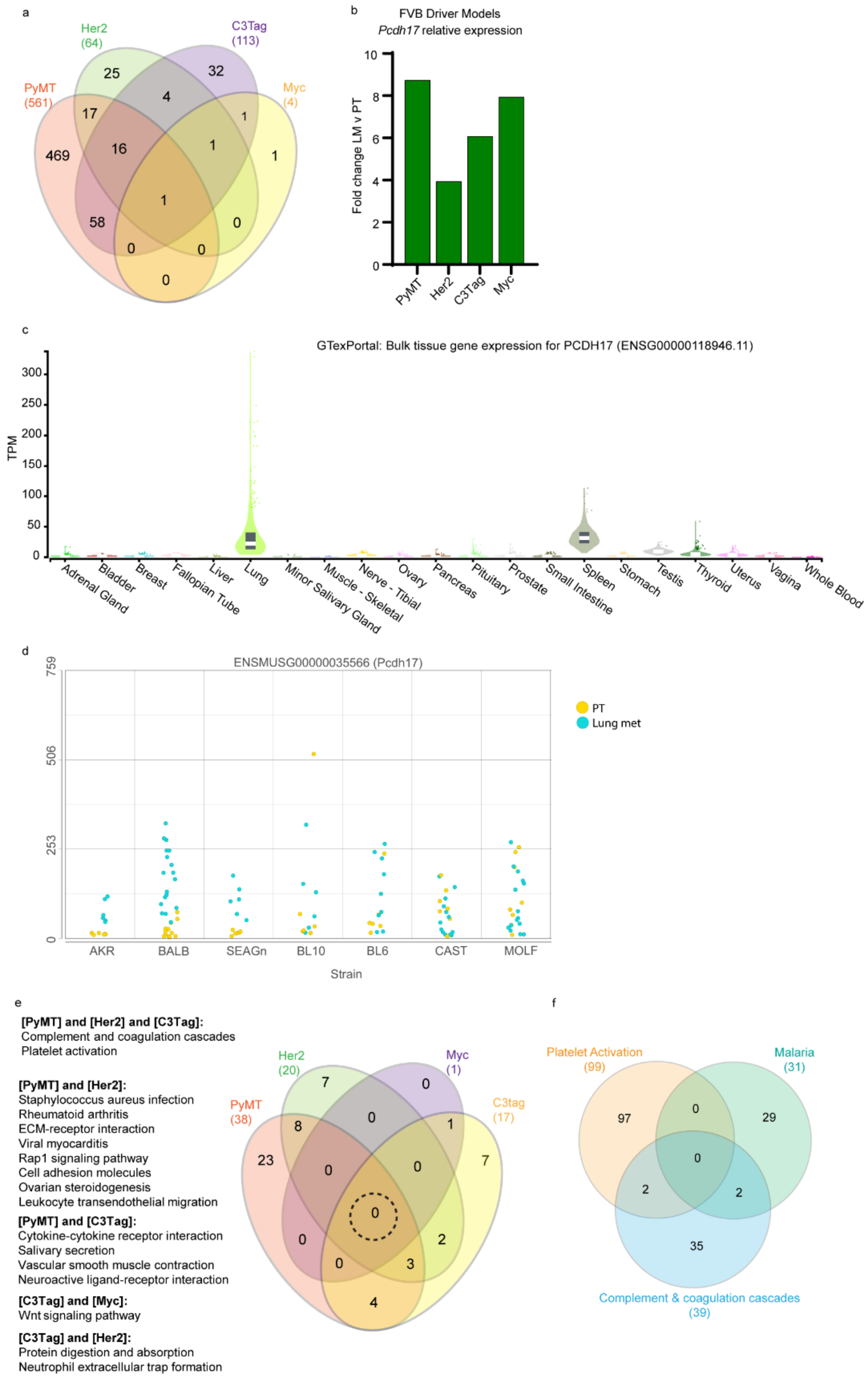
a
Her2
(64)
C3Tag
(113)
PyMT
(561)
Myc
(4)
25
32
4
17
1
16
1
469
1
1
58
0
0
0
0
b
FVB Driver Models
Pcdh17 relative expression
Fold change LM v PT
PyMT
Her2
C3Tag
Myc
c
GTexPortal: Bulk tissue gene expression for PCDH17 (ENSG00000118946.11)
TPM
Adrenal Gland
Bladder
Breast
Fallopian Tube
Liver
Lung
Minor Salivary Gland
Muscle - Skeletal
Nerve - Tibial
Ovary
Pancreas
Pituitary
Prostate
Small Intestine
Spleen
Stomach
Testis
Thyroid
Uterus
Vagina
Whole Blood
d
ENSMUSG00000035566 (Pcdh17)
PT
Lung met
AKR
BALB
SEAGn
BL10
BL6
CAST
MOLF
Strain
e
[PyMT] and [Her2] and [C3Tag]:
Complement and coagulation cascades
Platelet activation
[PyMT] and [Her2]:
Staphylococcus aureus infection
Rheumatoid arthritis
ECM-receptor interaction
Viral myocarditis
Rap1 signaling pathway
Cell adhesion molecules
Ovarian steroidogenesis
Leukocyte transendothelial migration
[PyMT] and [C3Tag]:
Cytokine-cytokine receptor interaction
Salivary secretion
Vascular smooth muscle contraction
Neuroactive ligand-receptor interaction
[C3Tag] and [Myc]:
Wnt signaling pathway
[C3Tag] and [Her2]:
Protein digestion and absorption
Neutrophil extracellular trap formation
Her2
(20)
Myc
(1)
PyMT
(38)
C3tag
(17)
f
Platelet Activation
(99)
Malaria
(31)
Complement & coagulation cascades
(39)

Supplemental Figure 8 MSGE in FVB driver models

(a) Venn diagram analysis of MSGE from each FVB driver model. (b) Histogram showing fold change in expression of *Pchd17* in LM vs. PT across FVB driver models. (c) GTex portal violin plots showing tissue-specific expression of human *PCHD17* (Transcript per million (TPM)). (d) Dot pot showing PyMT-F1 model expression of *Pchd17* where the y-axis shows normalized counts. (e) Venn diagram analysis of MSGE-enriched pathways for each FVB driver model. (f) Venn diagram analysis of gene lists assigned to Platelet activation, Malaria, and Complement and coagulation cascade pathways.

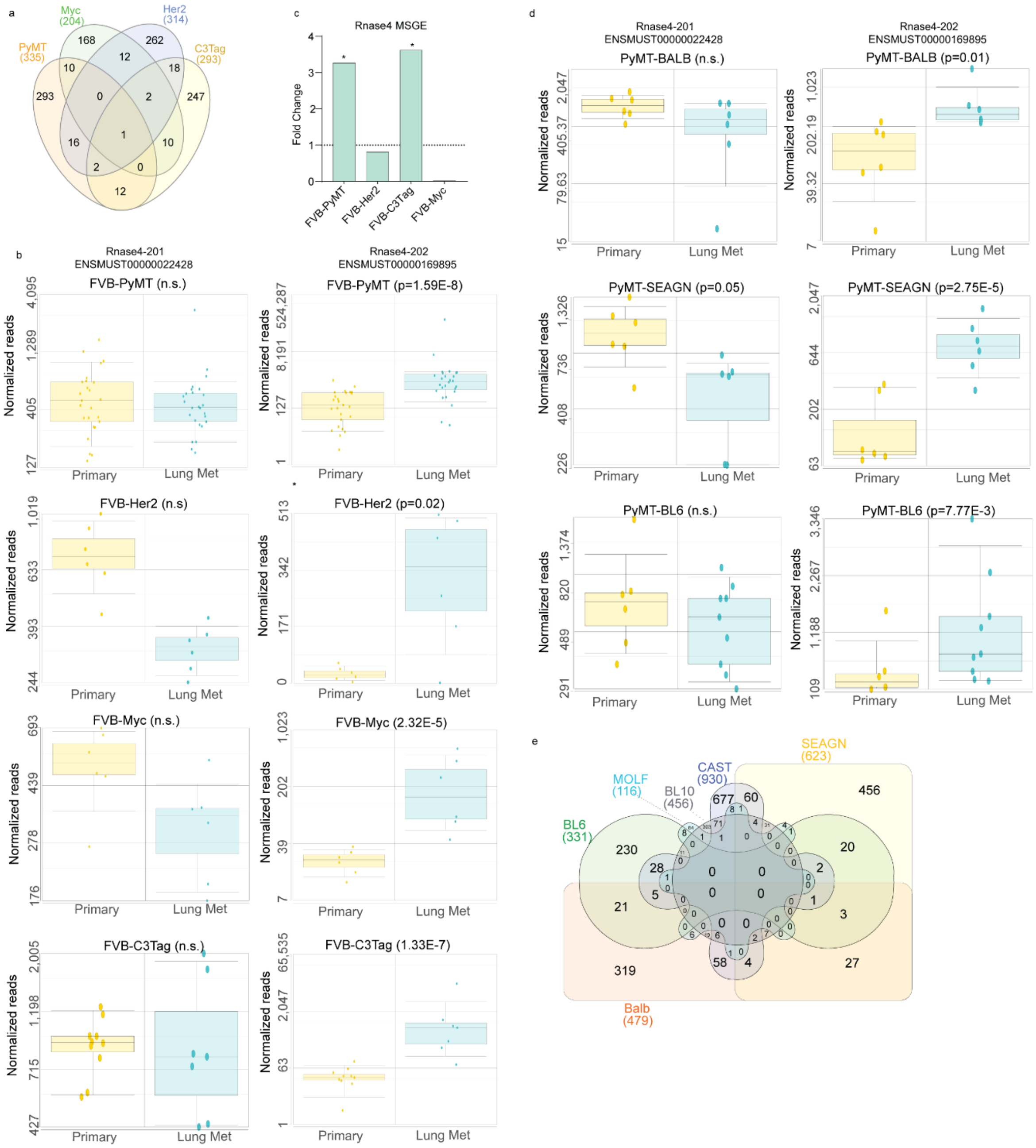

Supplemental Figure 9 RNase4 splice variant expression.

(a) Venn diagram analysis of transcript lists from LM vs. PT alternative splicing analysis in FVB driver models (p<0.05). (b) Paired box and whisker plots for *Rnase4-201* and *202* isoform levels in PT and LM tissue from each FVB driver model. (c) Histogram showing fold change in expression of the *Rnase4* gene in LM vs. PT across FVB driver models. (d) Paired box and whisker plots for *Rnase4-201* and *202* isoform levels in PT and LM tissue from BALB, SEAGN, and BL6 PyMT-F1 models. (e) Venn diagram analysis of transcript lists from LM vs. PT alternative splicing analysis in PyMT-F1 models (p<0.05).

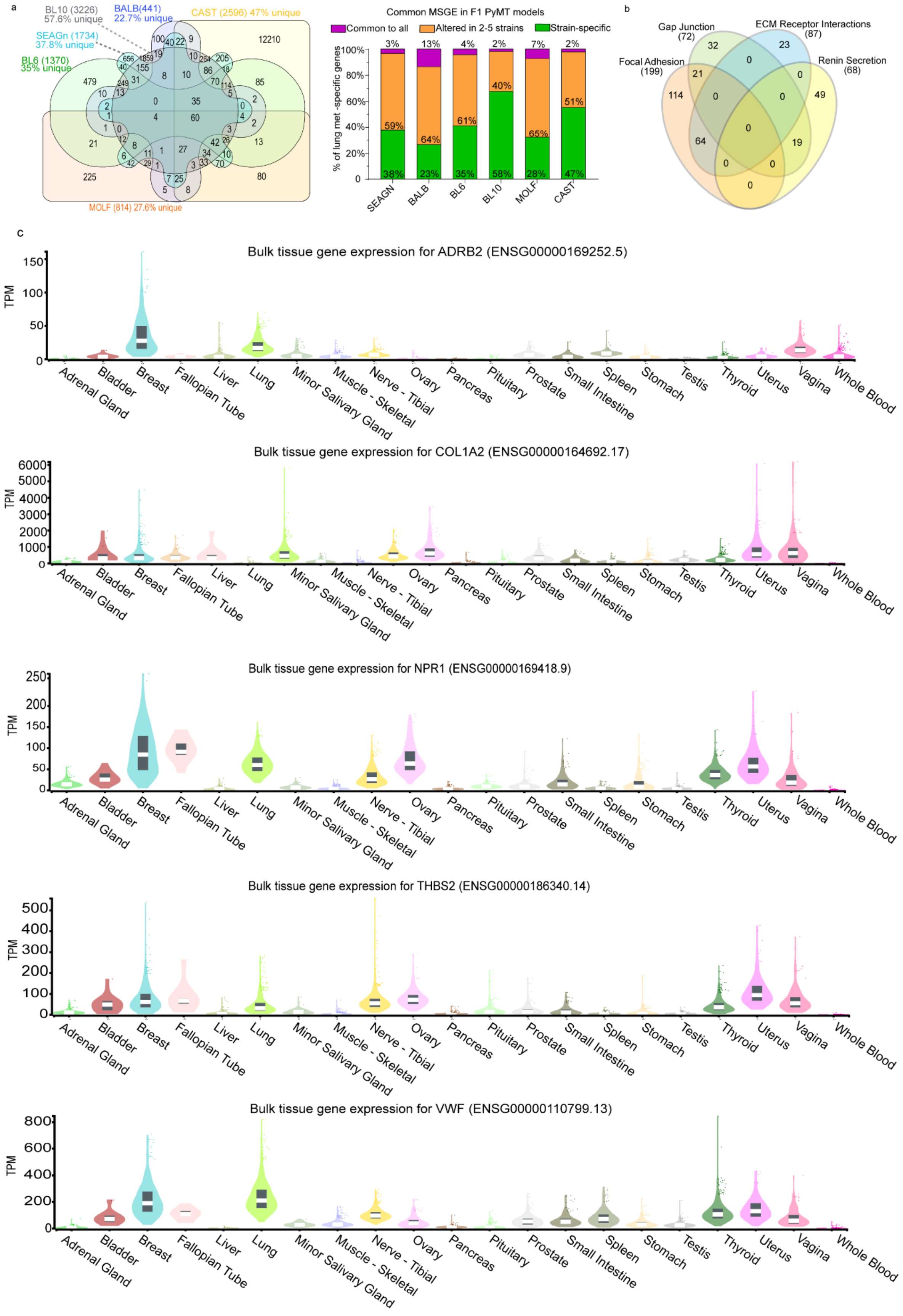
a
BL10 (3226)
57.6% unique
BALB(441)
22.7% unique
CAST (2596) 47% unique
SEAGn (1734)
37.8% unique
BL6 (1370)
35% unique
MOLF (814) 27.6% unique
Common MSGE in F1 PyMT models
Common to all
Altered in 2-5 strains
Strain-specific
% of lung met -specific genes
SEAGN
BALB
BL6
BL10
MOLF
CAST
b
Gap Junction
(72)
ECM Receptor Interactions
(87)
Focal Adhesion
(199)
Renin Secretion
(68)
c
Bulk tissue gene expression for ADRB2 (ENSG00000169252.5)
Bulk tissue gene expression for COL1A2 (ENSG00000164692.17)
Bulk tissue gene expression for NPR1 (ENSG00000169418.9)
Bulk tissue gene expression for THBS2 (ENSG00000186340.14)
Bulk tissue gene expression for VWF (ENSG00000110799.13)
TPM
Adrenal Gland
Bladder
Breast
Fallopian Tube
Liver
Lung
Minor Salivary Gland
Muscle - Skeletal
Nerve - Tibial
Ovary
Pancreas
Pituitary
Prostate
Small Intestine
Spleen
Stomach
Testis
Thyroid
Uterus
Vagina
Whole Blood

Supplemental Figure 10 PyMT F1models common MSGE

(a) Venn diagram analysis of MSGE from each PyMT-F1 model and a histogram summarizing the extent of gene overlaps within the comparison. (b) Venn diagram analysis of the gene lists assigned to Focal adhesion, Gap junction, ECM receptor interaction, and Renin secretion pathways. (c) GTex portal violin plots showing tissue-specific expression of *ADRB2*, *COL1A2*, *NPR1*, *THBS2*, and *VWF* (Transcript per million (TPM)).
